# Emergence of focal cortical hyperexcitability in the murine tetanus toxin epilepsy model

**DOI:** 10.64898/2025.12.31.697069

**Authors:** Jochen F Meyer, Stelios M Smirnakis

**Affiliations:** Baylor College of Medicine, Department of Neurology, Houston, TX 77007; Brigham and Women’s Hospital and Jamaica Plain Veteran Administration Hospital, Harvard Medical School, Boston, MA 02115

## Abstract

Focal epilepsies are a heterogeneous group of neurological disorders that may be caused by a large variety of underlying brain lesions. Long-term quality of life prognosis is often poor, and treatment options are limited. Despite significant progress in animal model sophistication over the past decades, a deeper understanding of epileptogenic processes on a mesoscale and microscale circuit level is needed. Here we model focal epileptogenesis in mice using single intracortical tetanus toxin injections and employ chronic in vivo two-photon and widefield calcium imaging with simultaneous EEG and behavioral video recordings (“multimodal” recordings) to follow epileptogenic changes over time. We observe a moderate increase in global brain excitability on EEG but, more strikingly, a gradual emergence of brief but intense optically recorded epileptiform events (“microseizures”) that are spatially limited, peaking around 20-30 days after toxin injection and waning over the next 30 days. During these events, a majority of local neurons are recruited into extreme depolarization, yet there is considerable variability in duration and spatial coverage between events, depending on the proximity to the injection focus and the number of days elapsed since injection. Notably, EEG electrodes <3 mm from the focus typically fail to detect these events, highlighting the utility of high-resolution in vivo optical imaging for capturing all events in focal epileptogenesis. Single-cell patch clamp recordings in awake animals confirm that microseizure episodes are associated with depolarization block, during which the calcium signal remains high. Using chronic multimodal recordings, we detect an initial rise of interictal activity in local cortical networks during the first day after injection, followed by a layer-specific suppression and dysregulation of interictal spontaneous network activity in local cortical circuits preceding microseizure events, and low-level suppression when microseizures started subsiding. The chronic suppression in interictal activity is accompanied by relative hypoactivity of PV+ interneurons and concurrent hyperactivity of SOM+ interneurons. This integrated imaging-electrophysiology approach provides a powerful platform for unraveling circuit-level mechanisms of focal epilepsy and potentially for identifying and testing novel therapeutic strategies in the future.

## Introduction

Focal epilepsy can be caused by a variety of underlying insults such as brain trauma, vascular lesions, benign and malignant neoplasms. Besides the emergence of electrographic and behavioral seizures, there have been numerous reports of other neurological impairments such as cognitive deficits or neuropsychiatric disease linked to focal injury or ischemia leading to subsequent focal hyperexcitability^1,2^. The persistent nature of refractory seizures constitutes a profound medical challenge, and the power of diagnostic and therapeutic methods to treat these disorders is currently limited both by diagnostic sensitivity and lack of target precision manifesting in frequent pharmacoresistance^3,4^. The pathogenesis of focal hyperexcitability leading to epileptogenesis remains obscure to date, and few studies exist that follow the evolution of local circuit hypersynchronization chronically using a suitable *in vivo* model that does not involve permanent cellular damage as a confounding byproduct, with the possible exception of early stage electrical kindling models^5^ , a PTEN-KO model^6^ (although changes in cellular morphology were reported), and an optogenetic model, which however requires several weeks of consistent stimulation to induce^7^. A more detailed understanding of the processes underlying focal epileptogenesis at a cellular level is needed.

Among a variety of animal models for focal epileptic syndromes, the tetanus toxin (TeNT) injection model in mammals has proven useful in modeling pathological neuronal synchronization in a variety of brain areas and under a number of dosing and injection regimens (reviewed in ^8,9^, see also ^10–12^ ), including as a model for the development of novel antiepileptic agents such as autoregulatory gene therapy^13^. It is generally thought that TeNT cleaves synaptobrevin (also known as vesicle-associated membrane protein, VAMP), an essential component of the SNARE complex^14,15^ that mediates presynaptic vesicle-to-membrane fusion, and binds to synaptic vesicle glycoprotein (SV2)^16^. Consequently, it inhibits release of neurotransmitters such as acetylcholine and glycine^17^ , as well as GABA, noradrenaline, and others^18,19^ . Near the injection site, TeNT causes a reorganization of spatiotemporal synaptic vesicle dynamics at inhibitory and excitatory synapses for 10-45 days after injection^20^, past the main proteolytic phase of toxin activity (10-21 days after injection^21^). The overall preferential blockade of inhibitory transmission^9,22–24^ causes excitation/inhibition imbalance locally that extends to nearby circuitry through diffusive processes, to generate epileptiform activity. However it is thought that the emergence of chronic epilepsy is mediated by pharmacoresistant, recurring seizures following TeNT injection, not by the toxin itself^8,10,12,25^. For example, Vannini et al^26^ showed that levels of intact VAMP2 in mouse cortex injected locally with TeNT are reduced for longer periods of time than TeNT proteolytic activity was present. Furthermore, when TeNT was injected in rat neocortex, GABA release was only impaired transiently by the toxin, but early primary and secondary epileptic foci appeared to cause long-lasting changes in local functional connectivity underlying long-term epilepsy^25^.

Most studies using TeNT as a model of focal, chronic hyperexcitability in mammals have focused on intra-hippocampal injections to study epileptogenic effects of TeNT ^11,27–30^. In cortex, this model has predominantly been studied in cats ^31^ and rats, ^10,25,32^ with a recent study suggesting changes in gene expression depending on the stage of epileptogenesis in the TeNT model^33^. Differences in toxin effects depending on the cortical area injected have been reported^34^. Yet, a critical gap remains in our understanding of how local circuit dysregulation within the targeted neuronal population translates into focal epileptiform activity and, ultimately, epileptogenesis in this model. Historically, such questions were difficult to address: cellular-resolution methods were restricted to patch-clamp recordings from only a few neurons at a time, while field potential techniques such as ECoG offered population-level activity with limited spatial precision. The advent of two-photon calcium imaging now provides the opportunity to bridge this gap by capturing activity patterns across large ensembles of neurons in vivo, offering new insights into how local network perturbations induced by TeNT evolve into pathological circuit states ^35^.

Here we employ in vivo longitudinal imaging to characterize the progression of TeNT-induced epileptogenesis in the mouse neocortex and to address several outstanding questions. For instance, it remains unclear how neuronal activity patterns at the single-cell level relate to the ‘epileptiform’ events detected on EEG. Likewise, we do not yet know how local ensembles of dozens to hundreds of neurons might develop gradually emerging abnormalities in circuit activation and organization that remain invisible to conventional EEG recordings. In addition, the dynamic changes in activity patterns that occur during the latent period—between the initial insult (TeNT injection) and the eventual emergence of overt epileptiform discharges—are still poorly understood. By imaging cortical circuits at cellular resolution over time, we seek to advance our understanding of how focal disruptions evolve into network-level pathology.

We used chronic EEG recordings together with simultaneous two-photon and widefield imaging of local cortical circuits in mouse V1 to study hyperexcitability development in the TeNT mouse model of chronic epilepsy arising from a cortical focus. First, we show that 5ng TeNT injection in mouse V1 induces abnormal cellular hyperexcitation events, most of which are invisible on EEG, highlighting the value of optical imaging as an adjunct to chronic EEG in studying epileptogenesis. In contrast, when we tested a substantially lower dose (0.15 ng), we found no evidence of significant network hyperexcitability, underscoring the importance of dose selection as suggested by prior studies^10^. Second, we delineate both transient and chronic changes in neuronal network activity patterns that evolve into epileptiform activity over time, focusing on brief, localized, high-amplitude network discharges which, to our knowledge, have not previously been described in the context of TeNT injections, though similar phenomena have been noted in other models of focal hyperexcitability^36^. Third, we define the spatial footprint of cortical network activity engaged by the TeNT injections using widefield one-photon imaging. Fourth, we probe the mechanisms of neuronal recruitment into seizures and epileptiform events by combining in vivo patch-clamping, two-photon calcium imaging, and EEG, and we characterize chronic changes in the relative activity of parvalbumin- and somatostatin-expressing interneurons. Together, these results suggest that the period leading to TeNT-induced emergence of microseizures , or spatiotemporally focal cortical network hyperexcitability events, is associated with inter-event activity suppression, a picture more nuanced than traditional excitation/inhibition imbalance models^37^. This argues for the use of new integrated approaches combining chronic EEG with longitudinal 2-photon imaging to dissect the circuit changes that occur during epileptogenesis, ultimately informing the development of improved strategies for early diagnosis and therapy of focal epileptic syndromes.

## Materials and Methods

All animal research was performed in accordance with the guidelines and regulations by the Institutional Animal Care and Use Committee (IACUC) of Baylor College of Medicine and VA Boston Healthcare System.

### Animal Surgery

Adult (4-6 months of age) male C57BL/6J-Tg(Thy1-GCaMP6s)_GP4.3Dkim/J mice (Jackson Laboratory, Stock Number 024275), or C57BL/6J PV-Cre/Ai9 (Jackson lines #017320, 007909), or C57BL/6J SST-Cre/Ai9 (Jackson lines #013044, 007909), or C57BL/6J-Ai96/Syn-Cre (Jackson lines #028866, 003966) were used for the experiments. See table 1 below for numbers and types of animals in the experimental groups (see also suppl. table 1 for more details).

**Table 1:**
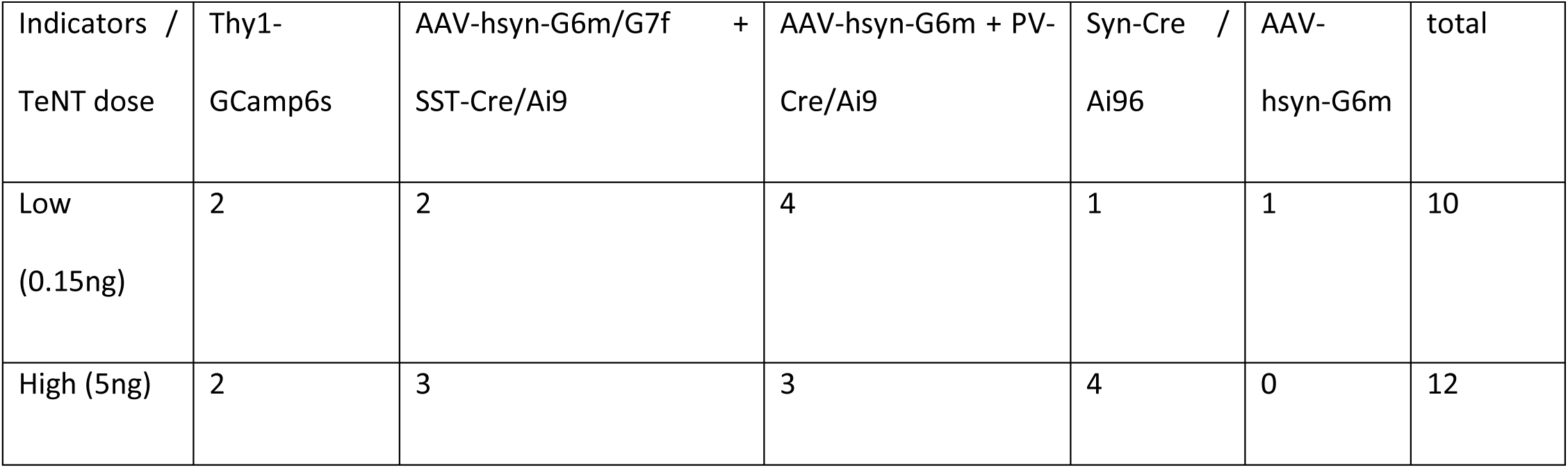
distribution of fluorescent reporter types across animals in this study.

Seven to fourteen days prior to TeNT or vehicle injection, animals were implanted with Electroencephalography (EEG) electrodes and a cranial window as previously described^38^ . Briefly, mice were anesthetized by inhalation of Isoflurane (4-5% in pure O_2_ for induction and 1% - 2% during surgery). EEG electrodes (Teflon coated silver wire with deinsulated, beveled and chlorided tips, diameter of 0.005 inch^39^) were inserted to the epidural space through cranial burr holes and secured by dental cement (Lang Dental, Wheeling, IL). Recording electrodes were placed bilaterally at 2.7 mm lateral and 3.5 mm rostral to lambda and reference electrodes were placed at 2 mm lateral and 2 mm posterior to lambda. A 3 mm diameter craniotomy (centered at 2.7 mm lateral and 1.5 mm rostral to lambda) was drilled and a coverslip was cemented in place with dental cement. A custom designed aluminum head post (emachineshop.com, Mahwah, NJ) was attached to the skull with dental cement for imaging.

### Tetanus toxin injection

Using a glass micropipette (tip diameter 30-50 mm), TeNT solution (Sigma-Aldrich Inc., USA, 200-800nl, 0.15 or 5 ng in phosphate buffered saline containing 0–2 % bovine serum albumin (BSA)) or saline+BSA (diluting solution without the TeNT) was injected to the visual cortex at a depth of 400-800 μm through a hole in the cover glass under 2-photon microscope guidance or through a burr hole next to the cover glass^12^. The depth of the injection was controlled by micromanipulator (MPC-200, Sutter Instrument Company, San Francisco, CA). The solution volume was controlled by an automatic nanoliter injector (Nanoject II, Drummond Scientific Company, Broomall, PA). By mixing in Alexa 594 dye (Sigma-Aldrich Inc.,USA) with the injection solution, we ensured, under direct 2-photon microscope visualization, that the solution spread evenly throughout all layers, though we cannot exclude that there might have been uneven toxin spread, resulting in a more concentrated delivery to L5/6 (see fig. 1b). Through visual inspection of the Alexa dye at low (4x) magnification, we estimate that the solution spread into a local volume of 0.8-1.5 µL on average. Because the presumed mechanism of action of TeNT is mediated primarily by functional suppression of GABAergic interneurons, we labeled these cells with tdTomato in a subset of animals using the PV-Cre/Ai9 and SST-Cre/Ai9 crosses to compare cell count and functional changes before and after TeNT injection.

**Figure 1.**
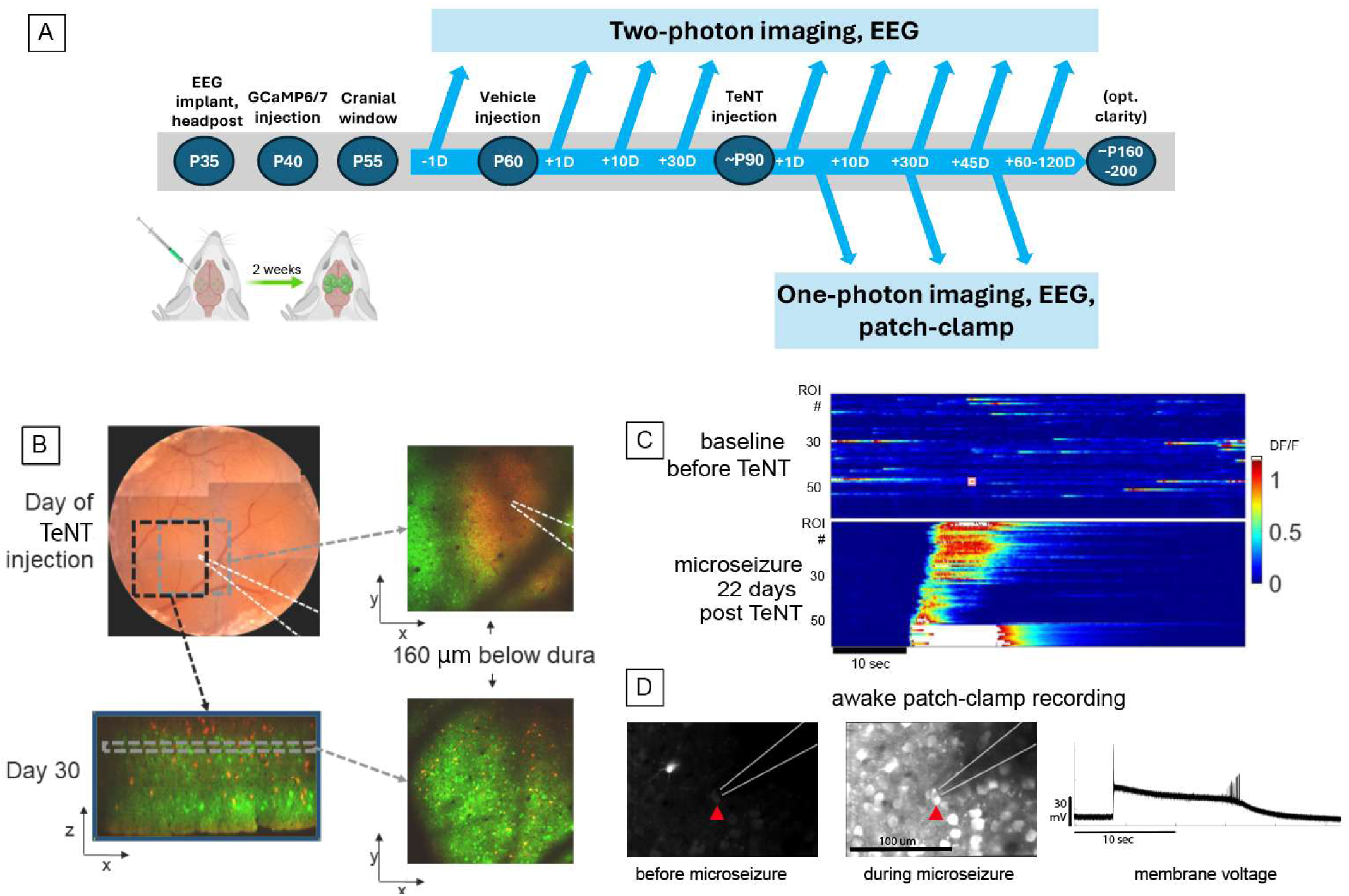
Experimental Paradigm. **A)** Mice received EEG wire implants and AAV-GcaMP6 (or GcaMP7) injections in primary visual cortex (V1) at 2-3 months of age, followed by a cranial window, vehicle injection and TeNT injection (see methods). There was at least one imaging session prior to toxin or vehicle injection, and after injection, imaging was carried out regularly for 60-120 days post injection. **B)** *<u>Top Row Left:</u>* Brightfield image of the 3mm diameter cranial window implanted. *<u>Top Row Right:</u>* Two-photon image of the same window at 160μm below dura, during TeNT injection (red = Alexa594 dissolved in the TeNT solution, green = GcaMP6). *Bottom Row Left*: Side view of the typical volume of neurons imaged, 30 days after TeNT injection (x/z projection from a 2-photon raster volume scan acquired from 0-600μm below dura. Green = GcaMP6, red = tdTomato expressed in PV+ interneurons. Bottom Row Right: x/y plane, 160μm below dura. **C)** Calcium activity traces from individual neuron regions of interest (ROIs) were extracted. *<u>Top:</u>* typical raster plot of a local group of neurons before TeNT injection; the last 10 rows represent calcium signal from extracellular neuropil patches. *<u>Bottom:</u>* raster plot from the same group of cells during a microseizure event, 22 days after TeNT injection. **D)** To analyze membrane potential changes during microseizure events, we performed awake single-cell patch clamp recordings. *Left Panel:* patched cell (red arrow) and surrounding neurons during quiet baseline activity, patch pipette outlined post-hoc. *Middle Panel:* patched cell and surrounding neurons during a microseizure event displaying high intensity calcium fluorescence. *Right Panel:* voltage trace from the patched neuron around a microseizure event showing strong depolarization throughout the microseizure. Total time shown = 27 sec.

We adopted the strategy of injecting with vehicle first, documenting that no significant change occurred acutely and over 1-4 weeks, and then injecting TeNT at a second, later time point. This has the benefit that within the same animal we had a pre- & post- vehicle injection baseline control. This was done in 1 low-dose animal, and 10 high-dose animals, with no significant effects of vehicle injections observed. For both the vehicle and TeNT injections we drilled a ∼0.5mm diameter hole in a 3mm coverslip, attached the coverslip with Vetbond only, injected using glass capillaries connected to a Nanoject II under two-photon guidance, and replaced the coverslip with an unmodified one afterwards, sealing it with Vetbond around the edges, and a thin layer of dental cement.

### Calcium imaging and EEG recording

Awake mice were imaged under a Prairie Ultima IV 2-photon microscope (Bruker, WI), head fixed by head posts attached to a custom-built imaging stage, which allowed the mice to run freely backwards and forwards on a wheel treadmill with minimal movement at the imaging site. We targeted L2/3 neurons (ranging 160 −250 μm under pia) using a 16X or 25X objective (0.8 or 1.1 NA, respectively, Nikon Instruments, NY, USA) under spiral scanning mode (10-12 Hz, typical FOV size 600 x 600 microns, 1.6-2 microseconds dwell time, laser power below 20 mW). For widefield recordings, cranial windows were imaged using a 4x objective (FOV 6×6mm). In 2 animals, widefield images were acquired using an sCMOS camera (pco Edge4.2, pco Germany) under one-photon excitation. EEG signals were recorded at 10 kHz (Prairie View) with a bandwidth of 0.1 Hz-1000 Hz and amplified (A-M System, Model 1700, WA). Before recording, animals were allowed to acclimate with the imaging stage for 20 minutes in darkness before the recording. EEG and/or Calcium signals were recorded with Prairie View software. Two or three 20 minutes episodes were recorded before (n = 9 low-dose + 12 high-dose animals) injection and at 1h (9 low-dose, 12 high-dose TeNT, 10 saline), day 1 (7 low-dose, 7 high-dose TeNT, 3 saline), days 2-20 (8 low-dose, 9 high-dose TeNT, 5 saline ), days 21-45 (4 low-dose, 12 high-dose TeNT and 5 saline), and beyond 45 days (up to 60-155 days post injection, depending on experimental factors such as stability of the head post assembly and cranial window; 3 low-dose, 11 high-dose TeNT) after injection.

Animal locomotion during the recording was monitored using a custom adapted angular velocity encoder. 17 of the animals were also recorded by IR wide-field video (camera DCC3240N, Thorlabs) at 20 fps to monitor behavior such as grooming and whisking. For each recording, periods of active running and whisking were excluded from further analysis of interictal EEG and calcium dynamics.

### EEG data analysis

EEG recordings were analyzed using Matlab (Mathworks, Inc, Natic, MA) with custom code partially following previously published parameters^12^. Briefly, EEG signals were downsampled to 500 Hz and filtered to remove DC signal components and 60 Hz noise. A semi-automatic algorithm identified segments of interest with significantly elevated relative power (>1.5*SD, min duration 1 sec) in the 1-30Hz band, and these segments were further classified by visual inspection. Epileptiform spikes were classified as single excursions of ∼30-500msec duration and >= 4X standard deviation (SD) amplitude above the baseline, similar to previously published methods^40^. Groups of spikes (interval < 1 sec) that lasted for more than 5 seconds and featured a stereotypical gradual increase and subsequent decrease in amplitude and frequency were picked as epileptiform generalized convulsive seizure activity. Spike-wave discharges (SWDs) featured a prominent peak in the 6-10Hz frequency band^41^. To account for fluctuating EEG baseline activity, the 1.5*SD threshold was updated for each EEG segment containing one automatically pre-identified abnormal event plus 3 seconds before and after the event. These sections were visually monitored to exclude abnormal recordings or motion artifacts. Uncategorized events (UC) were automatically detected segments that did not have a strong or long enough signature in the frequency band we use to detect SWD’s or convulsive seizures, and they consisted – unlike conventional interictal spikes – of a series of deflections, typically of varying amplitudes (see suppl. fig. 1 for an example and average event rate across animals). Artifacts due to motion or electrical noise were excluded for each predefined candidate event by visual inspection. See figure 2 and suppl. figure 1 for examples, and supplementary figure 5 for an example of the EEG analysis algorithm.

**Figure 2.**
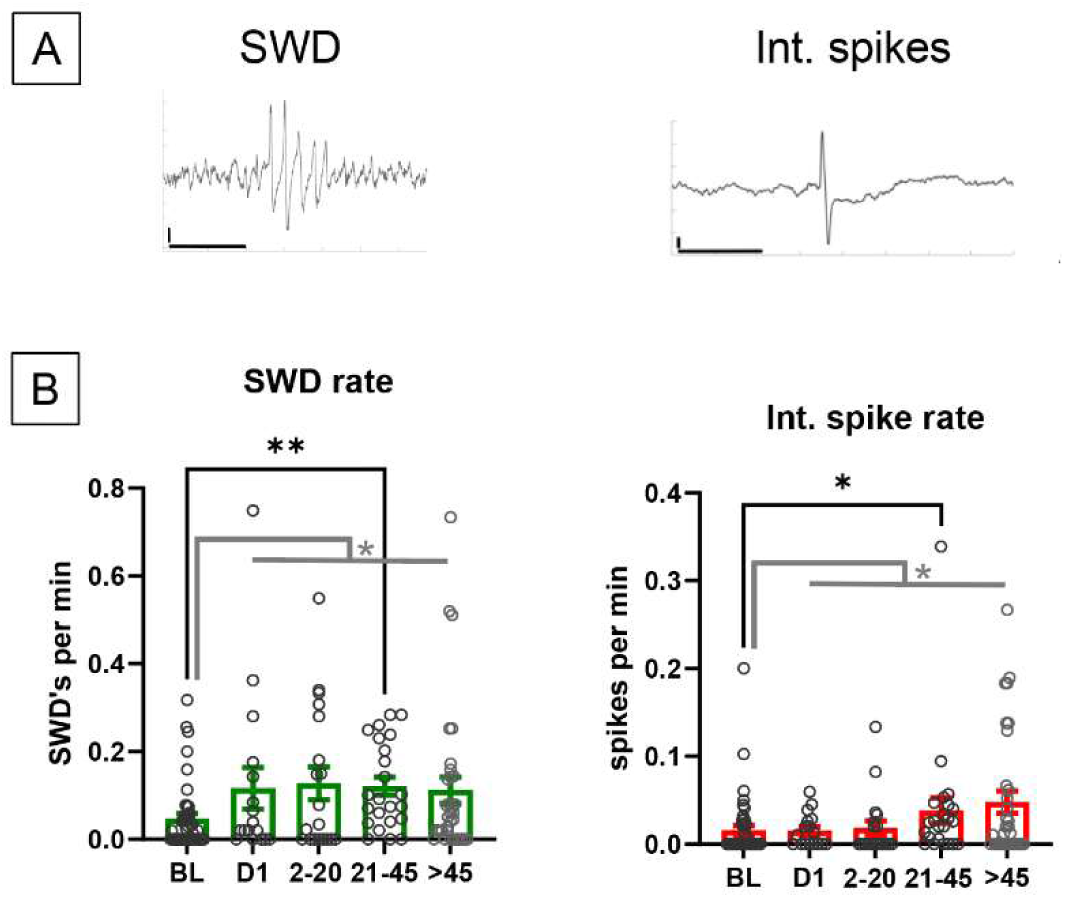
**Abnormal, epileptiform EEG events measured over time**: **A)** *Left:* typical example of a spike and wave discharge (SWD). Horizontal bar = 1 sec*<u>. Right:</u>* Illustration of an interictal spike. Horizontal bar = 1000 ms (left), 200 ms (right). Vertical bars = 0.1 mV **B)** *Left:* High-dose TeNT injection induced an increase of the rate of spike-wave discharge patterns seen on EEG compared to baseline. Comparing across individual time bins, the difference reaches significance (p = 0.009, Kruskal-Wallis test with multiple comparisons) on the bin corresponding to days 21-45 post injection *Right:* High-dose TeNT injection induced an increase of the rate of interictal spikes seen on EEG compared to baseline by the bin corresponding to days 21-45 post injection.

#### Calcium signal analysis

Calcium signal was analyzed based on previously published methods^38^ . Raw tiff image stacks were loaded into Matlab, motion corrected using normcorre, and constrained non-negative matrix factorization (CNMF)^42^ was applied to identify active neurons in the FOV. ΔF/F traces were computed for each neuron, followed by deconvolution and neuropil contamination correction (adapted from suite2p-wrapperdeconv function^43^) using an algorithm based on previously published work^44^, yielding a deconvolved ΔF/F matrix (termed “dΔF/F”) that was used for all subsequent analysis steps. This represents an estimate of the current probability that one or multiple spikes were produced within a time bin corresponding to the inverse of the scan rate (typically ∼10-12 Hz for spiral scan, and 15-30Hz for resonant scan). Therefore, bursts of spikes occurring within ∼33-100ms generally show up as one distinct “event”, as do single spikes. Next, whisking, wheel velocity, EEG and photodiode voltage traces were downsampled and aligned to the dΔF/F traces. Periods of active whisking and running were excluded for the remainder of the analysis. Individual deconvolved calcium transients were identified by thresholding at the median+2*std, and 8 metrics were extracted: 1) overall (averaged) dΔF/F per min, 2) events/sec, 3) amplitude of the transients, and 4) area under the curve (AUC) of the transients. The 5^th^ −8^th^ metrics we computed for each recording are derived from characterizing functional connectivity: To identify coactive clusters of neurons comprised of >2 neurons, the clusterONE algorithm (BrainConnectivityToolbox, matlab^45,46^) uses the previously computed pair-wise Person correlation coefficient matrix as its weighted network input, and identifies potentially overlapping groups of at least 3 associated neurons. Clustered neuron ensembles have previously been identified from neuronal calcium activity data using very similar approaches^47–49^. The weighted clustering coefficient of each neuron represents the tendency of its neighbors to be interconnected (see equation (9) in ^50^ :

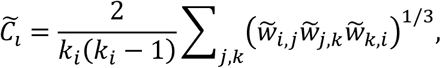

where k_i_ is the degree of a node, and 𝑤∼ are scaled weights of adjacent nodes.)

From this, we derived 6) the number of coactive clusters per area, 7) the overlap between clusters, and 8) density of nodes (i.e. neurons) within each cluster. For automated detection of microseizure events, we computed a synchrony (metric 8) trace for each recording corresponding to the fraction of coactive cells in each imaging frame. This trace was high pass filtered at 0.5 Hz, and any frames exceeding 6SD above the mean across the recording were considered microseizures. See supplementary figure 4 for examples of the microseizure detection algorithm as well as validation using shuffled distributions. Spatial and temporal extent of microseizures in widefield calcium sequences was identified by thresholding the ΔF/F signal at 7SD (chosen empirically for robust identification) above background fluorescence.

### Statistics

Non-parametric rank-sum Kruskall-Wallis test followed by Bonferroni’s correction for multiple comparisons was used for multi-group tests. For single comparisons of independent samples, the Wilcoxon rank-sum test was applied. For the comparison of chronic activity changes in 2-p calcium data, we used estimation statistics^51^. Estimation statistics, in contrast with hypothesis testing, determines significance by creating a bootstrapped 95% confidence interval through repeated (10,000 iterations) resampling with replacement. This allows for a direct overview and visualization of effect sizes when comparing paired values between the baseline group and consecutive time bin groups (figure 6). In these plots, comparisons are only shown when the 95% confidence interval of the paired differences (blue vertical line) lay completely above or below the zero line corresponding to no difference from the mean of the control group.

## Results

### Chronic 2-photon imaging enables long-term studies of focal TeNT epileptogenesis

Mice received a cranial window, vehicle, and/or TeNT injection without previous vehicle (see methods for details). We injected TeNT in layer 5 of primary visual cortex as previously described^12^. Prior to TeNT injection, 1 animal from the low-dose cohort, and 10 animals from the high-dose cohort were studied for 7-30 days after receiving vehicle-injections as controls, to differentiate the toxin-induced calcium and EEG patterns from naturally occurring neuronal activity that may exceed baseline characteristics in amplitude or frequency content. These animals did not develop any abnormal patterns of activity after vehicle injection (see supplementary figure 5 for a vehicle example).

There was at least one imaging session prior to toxin or vehicle injection and, following TeNT injection, imaging was carried out regularly until 60-120 days post toxin injection (figure 1a). We were able to follow neural activity in 22 animals for at least 10 days post TeNT injection, 18 of which were followed for >45 days. Imaging in all low-dose animals was restricted to layer 2/3 (100-300µm depth), while 6 of the 12 high-dose animals were imaged in both layer 2/3 and layer 4 (320-400 µm below the dura), the rest only in layer 2/3. Dual channel EEG was preprocessed (see methods) to remove artifacts, and abnormal (high power and amplitude; see methods) events were classified with a user-supervised semi-automatic algorithm (suppl. figure 5a) to identify episodes of spike-wave discharge (SW), seizures, interictal spikes, and other abnormal-appearing but uncategorized events (UC, see suppl. figure 5a for examples of UC and SW). TeNT was injected under 2-photon imaging guidance to verify the injection location and depth and volume injected (fig 1b). High-resolution z-stacks were acquired to examine morphological integrity of neurons during the post-injection phase, which we confirmed visually. No cell death was detected - and importantly, later during microseizure events the majority of neurons were highly depolarizable, demonstrating lack of permanent disability to modulate membrane potential.

Two-photon cellular calcium imaging allowed us to identify brief, recurring episodes of intense activity in which most neurons within the field of view (FOV) fired nearly synchronously, yet no concomitant deflection was evident on the simultaneously recorded EEG. Prior work has linked such synchronous local neuronal population calcium fluorescence events to intracortical LFP spikes in acute pro-convulsant 4-AP and pilocarpine models^36,52,53^, and combined optical–EEG recordings in the hippocampus of freely moving mice after kainic acid injections have captured optically defined epileptiform population transients that are visible on the EEG ^54,55^. Furthermore, “microseizures” confined to sub-millimeter domains have also been detected with high-density microelectrodes in rodents^56^ as well as with intracranial micro-electrodes in human cortex, where they were not detected on simultaneously recorded epidural surface electrodes^57^. Such events appear to be similar to the events we observe here. To our knowledge, this is the first demonstration in the TeNT model of such brief, exceptionally synchronous ensemble —hereafter ‘microseizures’— that emerge and recur over time after injection (Fig. 4a; three events within a continuous 30-min trace).

### Tetanus toxin injected locally in V1 rarely generates generalized tonic-clonic seizures, but other epileptiform events are seen on EEG over time

Although generalized tonic-clonic seizures (GTCS’s) were exceedingly rare (only observed in 3 out of 12 mice total), some EEG abnormalities were seen in the EEG of all animals injected with TeNT over time (see suppl. fig 1 for number of mice displaying the 4 different types of EEG discharges per time bin). This is likely because we performed EEG monitoring only under the 2-Photon microscope, thereby recording for a maximum of ∼24 hrs across all time points of observation. The frequency of TeNT induced seizures reported in the literature is between zero^21^ (TeNT in mouse V1) and ∼0.85 (TeNT in rat V1) convulsive seizures per day^13^ , so the fact that we observed GTCS’s on EEG for only 3 out of the 12 mice we recorded is not altogether surprising. However, since it is possible we missed a significant fraction of generalized seizures because of under-sampling, we did not analyze seizure characteristics any further. Instead, we extended prior work by systematically characterizing abnormal epileptiform EEG patterns and found that intra-cortical 5 ng TeNT injections result in spike-wave discharges (SWDs) and interictal spikes that increase in frequency gradually over time (figure 2a). Specifically, we observed a sustained increase in SWDs, by 154% percent between D20-45 post injection compared to baseline (p = 0.009, Kruskal-Wallis test with multiple comparisons, fig. 2b). By comparison, interictal spikes emerged somewhat more gradually until reaching significant levels at 20-45 days (p = 0.048, Kruskal-Wallis test with multiple comparisons, fig. 2b), with low baseline levels of spikes recorded beforehand, indicating that the two types of events may originate from at least partially independent underlying mechanisms^58–61^. Uncategorized (UC, see methods) events were seen in 12/12 animals (total), and even though we do not believe they represent artifacts, the variability of their occurrence precluded statistical significance between baseline and subsequent time bins after TeNT injection: When pooled together, they occurred after TeNT injection 2.6 (±0.26 sem) times/min, versus 2.3 (±0.39 sem, p = 0.154 wilcoxon ranksum test), whereas SWDs and interictal spikes did increase significantly between baseline and post injection timepoints (SWDs: 0.0476 ±0.01 sem / 0.12 ±0.016 sem, p = 0.004; Int. spikes: 0.016 ±0.005 sem / 0.033 ±0.006 sem, p = 0.02, wlicoxon ranksum test, fig. 2b, grey significance bars). Perhaps not unexpectedly, low (0.15ng) TeNT injections did not appear to cause any seizures nor change the frequency of any of the 4 epileptiform EEG event types over the course of our recordings (suppl figure 1b).

#### Widefield imaging reveals high-intensity microseizure events

Using our multimodal chronic recording technique that allows us to record EEG and image calcium activity at cellular resolution as well as in widefield, low-magnification mode in the same animals, we screened the imaging data for any abnormal activity patterns that emerged after 5ng TeNT injection. Specifically, we employed mesoscopic 1-photon widefield recordings to quantify the overall spatial extent and spread of microseizure events after TeNT injection. Using a fluorescence intensity threshold of 7 SD above the median for each pixel in each FOV, we reliably identified spatially contiguous microseizure events and computed their average max. diameter (1.34mm +/- 0.284 sem), average ΔF/F amplitude (129% +/- 12% sem), and average duration (0.3s ± 0.1 s sem): Microseizure events were variable in amplitude, spatial and temporal extent (figure 3A). Remarkably, even though these events had spatial extent as large as ∼2 mm, we did not find significant elevation or suppression in EEG voltage or power around the microseizure time points (figure 3A top panel; see also suppl. Fig 3). Note, however, that the ipsilateral EEG electrode was located ∼3mm frontolaterally of the center of the imaging window, and the contralateral electrode was ∼6-7mm away. We did not observe activation of areas remote from the focus, however due to the limited size of the cranial window (3mm diameter) we cannot completely rule out distal or contralateral activation outside the purview of the window. These observations indicate that rather than causing a consistent, moderate elevation of firing rates in the local neuronal population, over time, TeNT pushes the network into an unstable activity regime that allows for spontaneous, brief episodes of extremely high activity amplitudes without generalization into widespread seizure states across other brain areas. The widefield recordings shown here were acquired on average around D43 (±13 sem) after TeNT injection.

**Figure 3:**
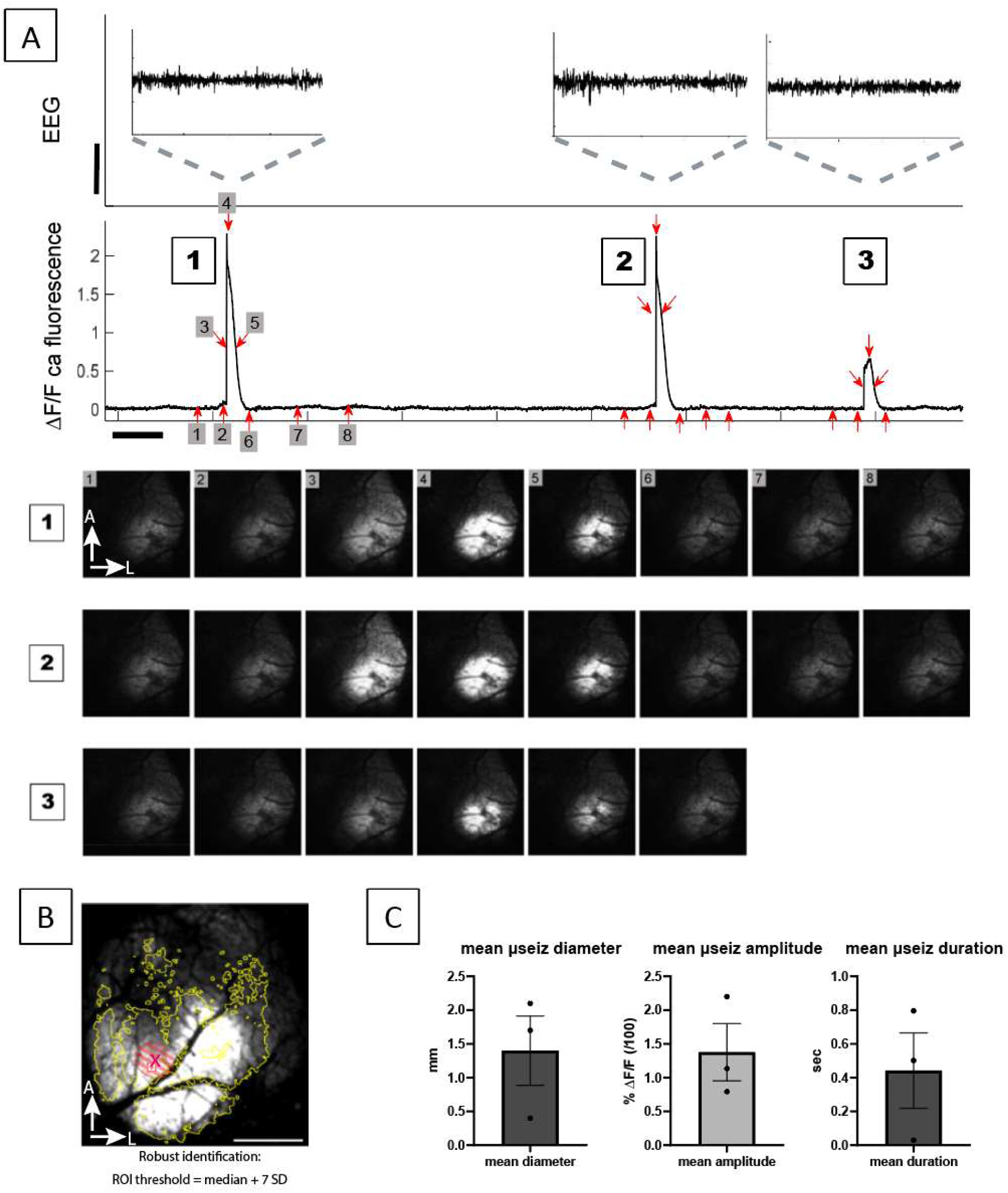
Widefield imaging outlines the spread of excitation patterns: **A)** Representative EEG and concurrently extracted mean calcium trace corresponding to several microseizures. The calcium signal is extracted from an ROI corresponding to a low-magnification microseizure (max. projection of all pixels around each microseizure time frames was averaged for all events in this recording to derive the ROI), measuring approximately 2mm in diameter. Horizontal scale bar = 30s, vertical bar = 0.5mV. This 8-minute recording segment contained three distinct microseizure events (1-3) of variable amplitude and duration (zoomed in insets above each microseizure for detailed visualization of the EEG). The snapshots below correspond to the time points marked with red arrows and numbers in the ΔF/F trace of calcium fluorescence above. Note that there is no well-defined evolution of the EEG signal corresponding to the optically defined microseizure event, recorded at an electrode that is ∼3mm away from the center of the window. **B)** Maximum intensity projection of calcium fluorescence during a microseizure event. Overlaid border (yellow) indicates the union of the individual ROIs derived from all microseizures observed in this recording, where each ROI was derived by thresholding each pixel’s intensity at 7 SD above the median. Dashed red disk shows the approximate extent of the initial spread of the TeNT injection visualized directly under 2-photon imaging (see methods). **C)** Bar plots (mean +/- sem) of average diameter (left), average amplitude (middle), and average duration (right), computed from 32 widefield microseizure events (3 animals, each data point corresponds to the mean per animal).

**Figure 4:**
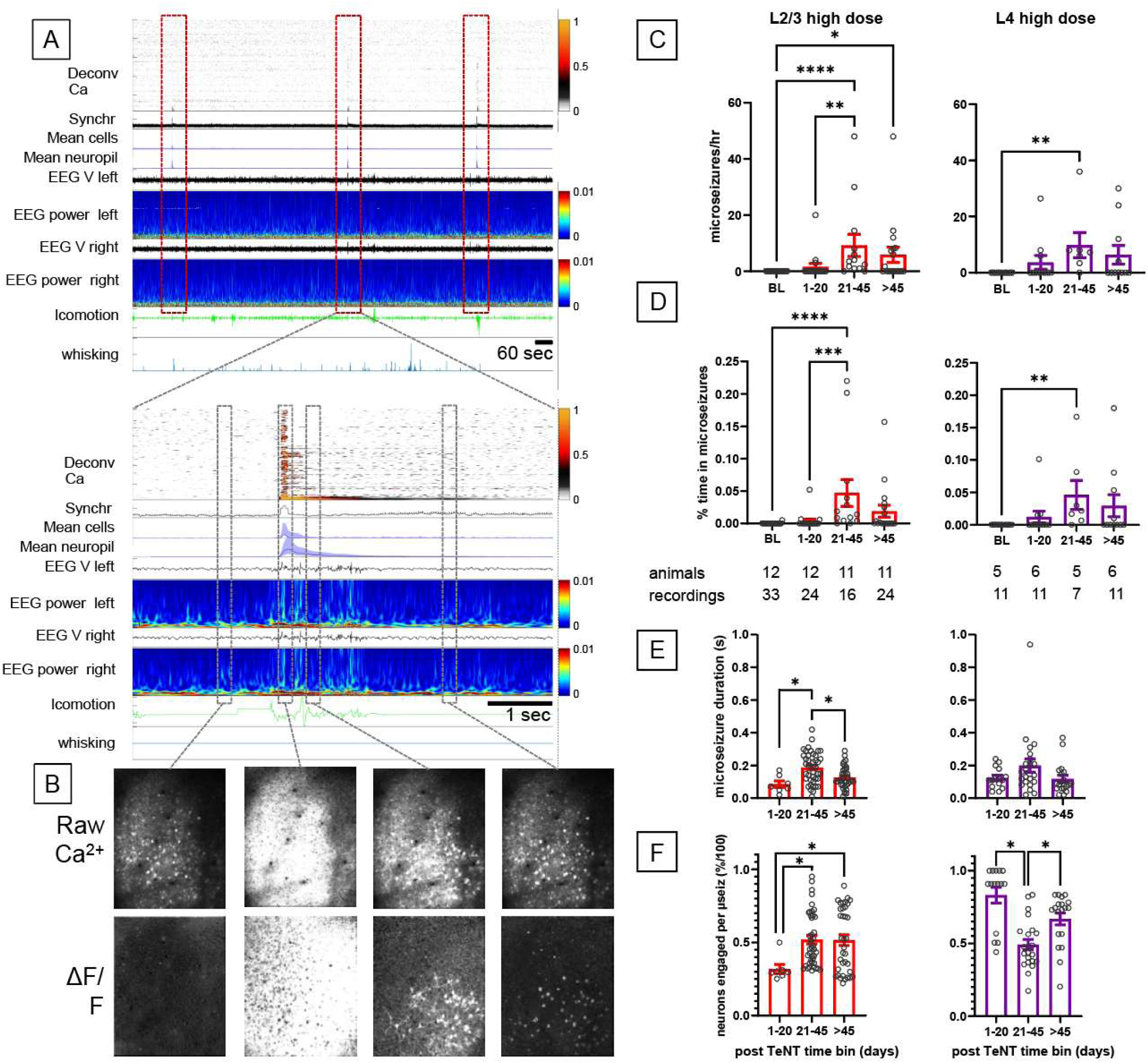
Time-binned analysis of microseizure parameters: **A)** *<u>Top:</u>* Multimodal 30 min recording featuring 3 individual microseizure events (red dashed boxes). *<u>Bottom:</u>* zoom in on one microseizure example from A, top. **B)** Snapshots of 2-photon acquired calcium activity at original frame rate 30 Hz, before, during and after the microseizure event from panel A. *<u>Top row:</u>* average of 3 frames (∼100 ms ) resonant scan raw fluorescence images. *<u>Bottom</u>*: after ΔF/F conversion. **C)** Number of microseizures per hour seen in L2/3 (left) and L4 (right) respectively, after injecting high (5ng) dose of TeNT. Bins reflect time interval: BL = baseline, days 1-20, 21-45, >45. **D)** Percent recording time spent in microseizures in L2/3 (left) and L4 (right), after a high TeNT dose. *<u>Below:</u>* number of animals and recording sessions used for generating the panels (C,D,E,F). Note that neurons in both layer 2/3 and layer 4 showed a significant increase in time spent in microseizure activity by days 21-45 after high-dose TeNT injection, with a subsequent reduction after day 45. Layer 4 neurons showed a tendency for time spent in microseizure activity to be increased earlier, from days 1-20, but this was not statistically significant. **E)** Mean microseizure event duration increased between early (D1-20) recordings and the time bin D21-45 (one data point per event is plotted), subsiding afterwards **F)** Mean fraction of engaged (synchronously active) neurons during microseizure events elicited after high dose TeNT injection. Note the early peak in neuronal participation from D1-20 in layer 4, compared to L2/3, where neuronal participation peaks later.

#### Microseizure intensity and frequency peak at 30 days, depend on distance from the injection focus

We used chronic calcium imaging to follow the activity of the same groups of neurons (with small fractions of cells possibly drifting in and out of the FOV’s across consecutive imaging time bins) over the duration of the recordings, up to 150 days post injection. To capture the behavioral state of the animal at all times, cellular calcium signals and EEG were supplemented with simultaneously recorded whisking and locomotion sequences (figure 4a). In order to compare microseizure fluorescence to baseline activity, we show snapshots of calcium activity before, during and after the microseizure example from fig 1C in fig 4A/B. Following TeNT injection, small groups of neurons started to develop hypersynchronous activity patterns that became more frequent over time (fig 4C/D). Abnormal calcium events first appeared around D10 in both cells and neuropil, increased in frequency and duration until D30-D45, then quieted down somewhat. Analogous to the EEG analysis, we quantified microseizure burden as events per hour (Fig. 4C) and as percent recording time (Fig. 4D), binned into four epochs: baseline (BL), 1–20, 21–45, and >45 days after TeNT injection (see methods). The high (5 ng) TeNT dose resulted in 9.2 ± 4 (sem) microseizures per hour in L2/3 and 9.8 ± 4.5 (sem) microseizures per hour in L4 recordings on average between 21-45 days post TeNT injection (fig. 4C). These were both significantly different than at baseline (BL vs 21-45: p 0.003 in L4 and p ≤ 0.001 in L2/3 respectively, Kruskal-Wallis test w/ MC). In L2/3 (fig. 4C), microseizure rates remained significantly different from baseline for periods >D45 (p = 0.019, all KW/mc test). Another way to assess the burden of microseizures is by plotting percent of time each cortical lamina is engaged in microseizures (fig. 4D). Again, both L4 and L2/3 showed significantly higher mean time spent in microseizures at 21-45 days compared to baseline (BL vs 21-45: p 0.0001 in L2/3 and p ≤ 0.0018 in L4 respectively, Kruskal-Wallis test w/ MC), with a trend towards higher values vs. BL persisting for D>45. Next, we quantified the mean duration of microseizure events after 5ng TeNT and found that in layer 4 recordings, microseizure duration was relatively consistent between time bins (0.12 ± 0.016 sem, 0.2 ± 0.041 sem, 0.118 ± 0.022 sem, respectively). In layer 2/3, microseizures were initially relatively brief, peaked around D21-45, and finally became shorter again after D45 (0.086 ± 0.02 sem, 0.187 ± 0.015 sem, 0.126 ± 0.01 sem, respectively; D1-20 vs. D21-45: p = 0.0084, D21-45 vs >D45: p = 0.01, Kruskal-Wallis test w/ MC, figure 4E). Interestingly, the mean fraction of neurons per FOV participating in microseizures evolved differently over time, depending on layer (figure 4F). In layer 4, participation peaked early at 83% ±5.4 (D1–20) then dropped to 49% ±3.6 (D21–45), with a partial rebound to 67% ±4.1 after D45. In layer 2/3, however, participation increased from 32% ±3 SEM on D1–20 to 52% ±3 on D21–45, and remained 52% ±4 after D45. This suggests that the temporal evolution of microseizure events likely begins in L4 and/or deeper layers closer to the center of the TeNT injection, and gradually spreads to encompass neurons in adjacent layers.

Mirroring the EEG event results (suppl fig 1), the 0.15ng TeNT dose, similar to controls, results in a very small number of microseizures in the early (D1-20) time bin that do not develop into more frequent events after day 21, resulting in no further microseizures observed in the D21-45 time bin, thereby failing to induce significant amounts of microseizure activity detectable by Calcium 2-P imaging (suppl. Fig 3).

#### Neurons recruited into microseizure events experience extreme membrane voltage depolarization

To elucidate membrane voltage changes underlying the single-unit activity observed in calcium imaging during microseizures, we performed in vivo whole-cell patch-clamp recordings in a subset of animals, simultaneously with EEG and two-photon calcium imaging. This approach enabled us to link single-cell intracellular membrane voltage with network-level excitability. Neurons patched within the microseizure focus, as indicated by high-intensity calcium fluorescence, exhibited prolonged membrane depolarizations that reached levels sufficient to block action potential generation for most of the microseizure duration (Fig. 5A–B). Despite this depolarization block, calcium fluorescence increased markedly (fig. 5B), reflecting sustained intracellular calcium accumulation rather than spiking activity.

**Figure 5:**
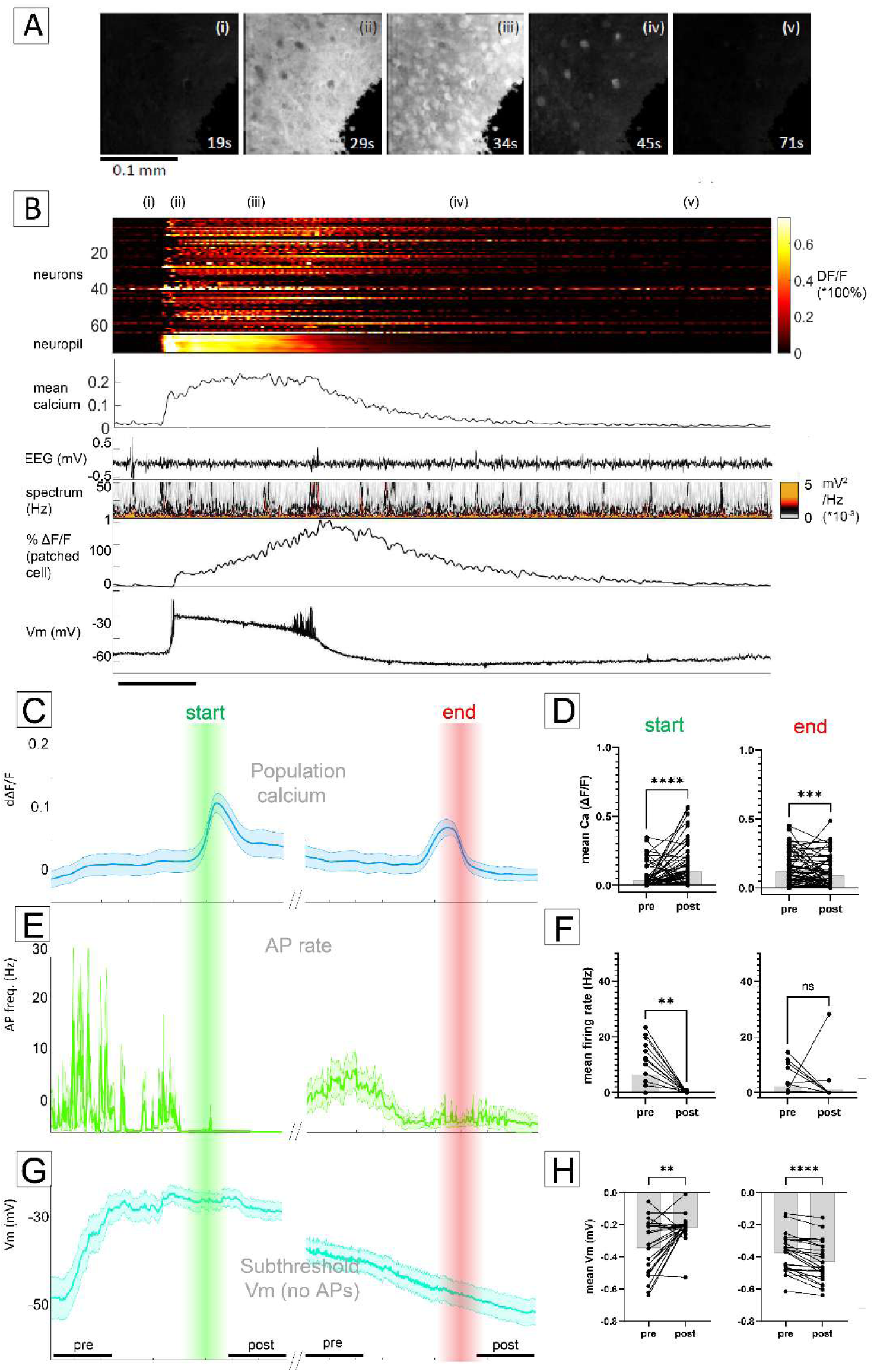
In vivo patch-clamp recordings verify and contextualize calcium imaging findings: **A)** Snapshots of an example FOV showing a microseizure event scanned at 10 Hz, while recording a single cell in current-clamp mode. **B)** calcium (% ΔF/F) traces extracted from the neurons in the FOV shown in (A), ipsilateral EEG voltage, EEG power, ΔF/F trace of the patched cell, and single-cell membrane voltage recorded during one microseizure event corresponding to the snapshots in (A). Start/end times were determined by thresholding the mean population calcium signal at mean + 6SD (methods) **C)** Calcium ΔF/F average (±s.e.m.) signal from all population averages in the 13 recordings (13 patched neurons, 6 animals, 1-5 microseizures per recording), aligned to the microseizure onset (green line) or offset (red line). **D)** Mean calcium ΔF/F activity across all patched units was compared before (average of a 1-sec interval starting 2.5 sec before and ending 1.5 sec before microseizure onset) vs. immediately (0 to 1-sec) after the microseizure offset. Groups were compared with a Wilcoxon rank-sum test. **E)** Action potentials extracted from patch clamp recordings (average of 13 cells from 6 animals) were used to compute aggregate single-cell firing rates aligned to microseizure onset and offset. **F)** Paired comparisons of firing rates before and after microseizure onset and offset, analogous to (D). **G)** After removing action potentials, membrane voltage (Vm) was averaged across the recorded neurons and aligned to microseizure onset and offset. Note the sharp >20mV depolarization in membrane potential 1-2 seconds before the microseizure onset versus the much more gradual repolarization across its offset. *Scale Bars: 1 sec* (across which averages were calculated) **H)** Analogous to (F), membrane voltage (Vm) increased significantly prior to microseizure onset and declined slowly around its offset. Microseizure onset/offset was defined by the corresponding rise/drop in population calcium activity measured across the respective FOVs.

**Figure 6:**
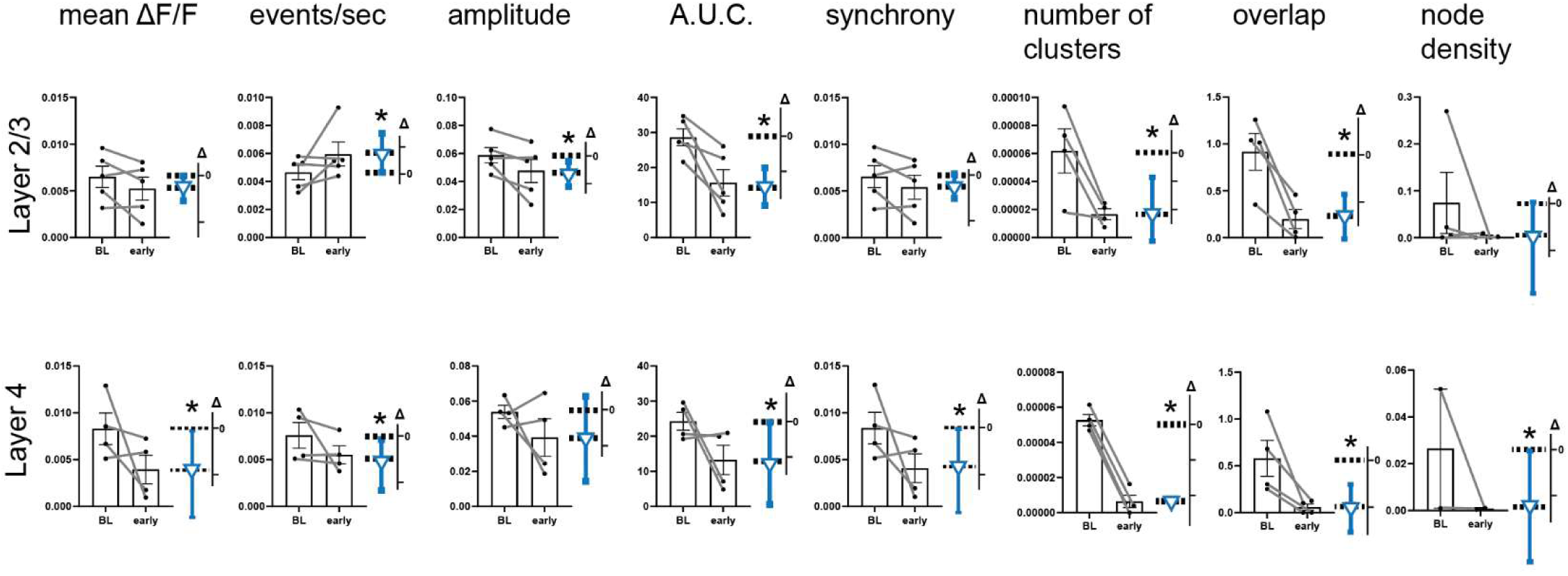
Changes in somatic activity and functional clustering over time, 1-20 days after TeNT injection. Each panel shows paired comparisons between baseline (BL) and post-TeNT measurements for the indicated metrics. Horizontal dashed lines represent mean values for the BL (equal to zero Δ) and early time bins. Blue vertical bars denote the mean difference ±95% confidence interval (CI) of the paired differences Δ (secondary y-axis units are the same as the primary axis’). CIs that do not intersect the zero Δ line indicate significant changes from baseline. During the early period (days 1–20), cellular activity measures in both layers 2/3 and 4 were significantly suppressed. Significant differences between BL and early time bins were observed for: mean ΔF/F, L4: difference = −0.0044, confidence interval (CI)[−0.0093 to −2.8e-4]; event rate, L2/3: diff = +0.0013, CI[−5.9e-6 to 0.0028]; L4: diff = −0.0021, CI[−0.0051 to −2.1e-4]; event amplitude, L2/3: diff = −0.01, CI[−0.021 to −0.0034]; AUC, L2/3: diff = −13, CI[−18.4 to −7.6], L4: diff = −11, CI[−22.6 to 0.54]; pair-wise synchrony, L4: diff = −0.0043, CI[−0.0093 to −2.2e-4]; clusters, L2/3: diff = −4.5e-5, CI[−6.5e-5 to −1.8e-5], L4: diff = −4.6e-5, CI[−4.8e-5 to −4.5e-5]; cluster overlap, L2/3: diff = −0.72, CI[−0.97 to −0.46], L4: diff = −0.52, CI[−0.81 to −0.28]; node density, L4: diff = −0.026, CI[−0.051 to −6.1e-4].

To quantify the temporal dynamics, we aligned ΔF/F fluorescence (Fig. 5C), spike rate (Fig. 5E), and intracellular membrane voltage of the patched cells (Fig. 5G) to microseizure onset and offset times, defined by the population calcium signal (Methods; Fig. 5B). Spike frequency, calculated from isolated action potentials, decreased sharply after microseizure onset (Fig. 5E-F), consistent with depolarization block occurring within several hundreds of milliseconds. No significant difference was observed around microseizure termination (fig. 5E-F), indicating that normal firing resumed and the depolarization block ended, before calcium fluorescence had decayed to baseline. After removal of action potentials, the mean membrane potential trace revealed that a sharp depolarization occurs approximately 1-2 seconds prior to microseizure onset, while a much more gradual repolarization towards the baseline is seen near the offset (Fig. 5G-H). Collectively, these data demonstrate that neurons participating in microseizures undergo intense and prolonged depolarization that suppresses spiking yet drives elevated calcium fluorescence, confirming that two-photon–detected microseizure events reflect *aberrant epochs of intense local network excitation*.

### Network dynamics leading to microseizures differ across layers

We next examined how local network activity evolved during the emergence of microseizures. We recorded calcium activity during periods of quiet wakefulness, isolated time periods that did not contain microseizure activity to measure only “interictal” patterns, and eight cellular activity metrics were extracted (mean dΔF/F, event rate, event amplitude, event area under the curve [AUC], mean pairwise synchrony, number of neuronal clusters per area, cluster overlap, and nodes per cluster), together with four corresponding neuropil patch measures (mean ΔF/F, event rate, amplitude, and AUC; Supplementary Fig. 6B) to characterize changes in network dynamics over time. Deconvolved calcium events were used to identify individual events corresponding to bursts of action potentials and higher-order co-active ensembles of neurons, using a previously published algorithm we adapted for our study^62^, and to compute and characterize changes in clustering parameters over time (Methods). Statistical comparisons were performed using estimation statistics^51^, contrasting baseline with immediate (≤2hr; Suppl. Fig. 7), early (days 1–20; Fig. 6), and late (days 21–45; Suppl. Fig. 6A) post-TeNT epochs. Plots in figure 6 compare pairs of time points, with average values from the same animals linked by a line. The vertical blue bar on the right represents the 95% confidence interval around the mean of the paired differences, while the horizontal lines are the means of the TeNT and the control (baseline) groups.

*<u>Immediate effects of TeNT:</u>* Suppl. Figure 7 illustrates changes observed within 2 hours after TeNT injection, thus capturing acute toxin effects. Overall, five nanogram TeNT had a modest effect within the first 2 hours after injection: mean neuronal calcium event rate increased in layer 2/3, accompanied by both higher event amplitudes and higher pairwise event synchrony. Event rate in layer 4 was also elevated (Suppl. fig. 7). These early changes are consistent with prior reports arguing that TeNT first preferentially inhibits GABAergic release, before causing broader, inhibitory and excitatory, synaptic disruption^63^.

*<u>Early phase (days 1–20):</u>* During the early post-injection phase—when microseizure activity first appeared—cellular and network-level properties changed across both layers 4 and 2/3 (Fig. 6). In layer 4, mean dΔF/F was reduced by 52 % (baseline: 0.008 ± 0.0017 s.e.m.; D1-20: 0.004 ± 0.0015 s.e.m.), driven by lower event rate and smaller event AUCs. Functional clustering, overlap between clusters and pairwise synchrony were also markedly suppressed, indicating loss of coordinated population activity. In layer 2/3, event amplitude and AUC were equally suppressed, however event rate was increased, suggesting a shift toward more frequent but weaker transients. Functional clustering and overlap between clusters were modestly reduced (fig. 6).

*<u>Peak Microseizure Epoch (days 21-45):</u>* After microseizure frequency peaked (D21-45; fig. 4C-D), activity metrics in layer 2/3 largely recovered to baseline (Suppl. Fig. 6, D21-45), whereas suppression persisted in layer 4 as reflected by decreases in mean dΔF/F, event amplitude, AUC, and synchrony (Suppl. Fig. 6). Node density (number of nodes per cluster divided by the cluster area), however, in layer 4 rebounded, reaching now above baseline levels.

*<u>Evolution of Neuropil Activity</u>:* Analysis of neuropil signals from small patches of neuropil adjacent to the neurons in the same recordings (10 patches per recording) revealed persistent laminar asymmetry. In layer 2/3, mean neuropil ΔF/F, amplitude, and AUC were significantly reduced compared to baseline throughout both early (D1-20) and late (D21-45) periods, whereas layer 4 neuropil activity did not significantly differ from baseline throughout (suppl. Fig. 6). Because neuropil fluorescence generally reflects aggregate axonal and dendritic signals akin to a high-resolution local field potential^64^, these results suggest that synaptic activity suppression in layer 2/3 persists at least to day 45 post injection, even though somatic AP activity metrics have returned to baseline by that time. Conversely in layer 4, neuropil activity metrics remain similar to baseline, even though cellular activity metrics show suppression, including at later time points (D21-45, suppl. figure 6A), when microseizures have peaked. These observations suggest that overall synaptic activity appears to be partially decoupled from somatic outputs.

Together, these findings reveal a temporally and spatially structured sequence of network changes during microseizure development. A brief, TeNT-induced increase in excitability is followed by a ∼20-day period of reduced interictal activity and disrupted clustering coinciding with microseizure onset. By days 21–45, layer 2/3 activity normalizes, whereas layer 4 remains partially suppressed. The persistent suppression of synaptic-level neuropil signals in layer 2/3, despite recovery of somatic firing, and conversely the persistent suppression of somatic signals in layer 4 while synaptic-level neuropil signal remain unchanged, points to a partial decoupling of synaptic input and spiking output—likely reflecting laminar differences in inhibitory–excitatory balance and connectivity^65–67^.

#### PV and SST-expressing interneurons in L2/3 are differentially modulated following TeNT injection

Previous studies have demonstrated that local TeNT injections induce disinhibition followed by the gradual emergence of epileptiform activity over several weeks^8^. However, this was mostly studied in the hippocampus^29,68^, leaving the specific contributions of cortical inhibitory and excitatory subpopulations to TeNT-induced disinhibition after intracortical injections incompletely understood^15,16^. To investigate how the two major classes of cortical inhibitory interneurons—parvalbumin-positive (PV⁺) and somatostatin-positive (SST⁺) cells—contribute to the emergence of cortical microseizures after TeNT injections, we used cell-type–specific labeling and longitudinal calcium imaging. In three high-dose animals, PV+ interneurons were labeled with tdTomato, and in three others, SST^+^ interneurons were labeled, to assess their anatomical integrity and functional differences as a function of their GABAergic nature. This was done by crossing PV-Cre or SST-Cre driver lines with a Cre-dependent tdTomato reporter (Ai9), yielding selective red fluorescence in each population (Methods). We did not observe anatomical changes in these cells across serial recordings. These interneurons, like the surrounding local neurons, also expressed GCaMP6. Representative examples from different time points before and after TeNT injection are shown in Fig. 7A,B. Neuronal densities of both PV⁺ and SST⁺ populations did not change significantly over time (Fig. 7C), indicating preserved anatomical integrity and lack of cell loss in the cortical region exhibiting microseizure activity, consistent with previous reports^31,69^. The apparent difference between SST^+^ density at baseline versus all later time bins is not significant, and importantly this apparent trend does not indicate a loss of SST^+^ interneurons, but rather a slightly higher density, which may be due to the red fluorescent reporter being more easily visible after residual obscurities of the cranial window disappeared over time.

**Figure 7:**
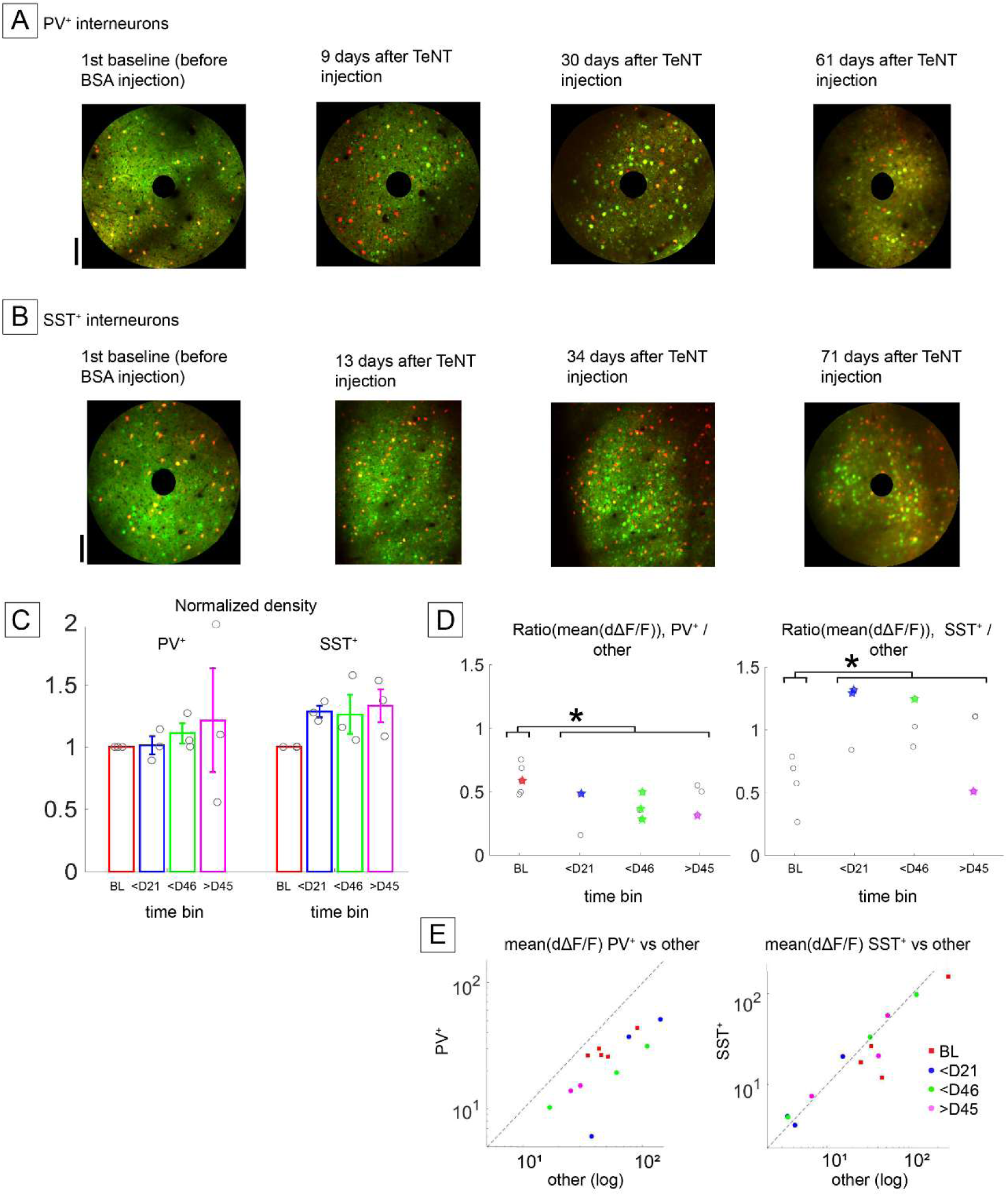
PV^+^ and SST^+^ population activity over time after TeNT injections. **A)** Dual-color overlay images from a representative animal expressing tdTomato in PV^+^ interneurons (red) and GCaMP6m in all neurons (green). The left image was taken before TeNT injection, followed by images at 9, 30 and 61 days after TeNT injection respectively. **B)** Analogous to (A), dual color overlay images from an example animal expressing tdTomato in SST^+^ interneurons, at −13, +13, +34 and +71 days relative to TeNT injection. Even though the images are not exactly identical they allow us to compute and follow the average density of interneurons over time. **C)** Quantification of PV^+^ and SST^+^ cell density, normalized by baseline recordings for each group. Groups were compared using nonparametric Kruskal-Wallis test with no significant results, indicating anatomical integrity of PV^+^ and SST^+^ cell bodies was not significantly impacted by TeNT injection. **D)***<u>Left:</u>* Quantification of changes in calcium activity levels between PV^+^ interneurons and putative pyramidal (“other”) neurons in the FOV. Each data point (gray circle) represents the ratio between the mean deconvolved calcium activity across all PV^+^ cells divided by the mean deconvolved calcium activity across all other cells in the FOV (one FOV per animal at each time bin). *<u>Right:</u>* Similar plot for the SST^+^ interneurons versus other neurons in the corresponding FOVs. For both panels, all baseline (BL) recordings were compared to all later recordings pooled together and found to be significantly different (Wilcoxon ranksum test, p = 0.045 for PV^+^/other comparison, p = 0.02 for SST^+^/other comparison). Colored stars denote recordings in which the calcium activity distributions of PV^+^ or SST^+^ cells were deemed significantly different from the calcium distributions of putative pyramidal cells in the same FOV. **E)** Scatter plots displaying the mean deconvolved calcium activity of putative pyramidal (“other”) neurons (x-axis) versus PV^+^ (y-axis, left panel) or SST^+^ neurons (right panel). Red squares: baseline; blue dots: mean activity corresponding to the early time period (D1-20); green dots: D21-45; magenta dots: >D45.

Although their numbers remained stable, activity levels of PV^+^ and SST^+^ interneurons changed over time following TeNT injection. Figure 7D shows the activity of PV^+^ and SST^+^ interneurons respectively. For each recording, we computed the mean dΔF/F for PV⁺ or SST⁺ cells, normalized to the mean activity of all other active neurons within the same FOV (predominantly pyramidal cells). These normalized activity ratios (PV⁺/other and SST⁺/other) were then plotted and compared across time bins (Fig. 7D). Different trends emerge for PV+ versus SST+ cells. PV⁺ interneurons showed a significant reduction in relative activity after TeNT injection (pooled over all post-TeNT time points) compared with the pre-TeNT injection baseline (p = 0.045, wilcoxon ranksum test; fig. 7D). In contrast, SST⁺ interneurons exhibited a significant relative increase for the same comparison (p = 0.02, wilcoxon ranksum test; fig. 7D). Scatter plots of the mean PV⁺ and mean SST⁺ neuron activity versus the activity of neighboring “other” neurons, across all animals further illustrate these opposing trends (Fig. 7E). Fig. 7E_left shows scatter plots of mean PV+ interneuron activity (y-axis) as a function of the mean activity of nearby putative pyramidal neurons (x-axis; “other” neurons in the same FOV), showing that most points (squares not colored red) are shifted towards lower values - further below the diagonal - compared to points obtained at baseline (red squares). In contrast, consistent with fig. 7D_right, scatter plot points for SST^+^ interneurons following TeNT injections are squarely on the diagonal, clearly corresponding to higher values compared to points (red squares) obtained at baseline (fig. 7E_right). These results suggest that, on average, in relation to neighboring putative pyramidal neurons, PV^+^ interneurons exhibit relative suppression compared to baseline following TeNT injections, whereas SST^+^ interneurons display correspondingly a relative increase in activity. Given that SST^+^ interneurons are known to inhibit PV^+^ interneurons, it is plausible that these dual effects — reduced PV-mediated inhibition and enhanced SST activity — may act synergistically to destabilize excitation–inhibition balance, weakening inhibitory control and promoting microseizure generation.

## Discussion

Focal epilepsies remain difficult to treat due to their multifactorial and frequently opaque circuit etiologies. This complexity, together with the absence of animal models that faithfully reproduce the chronic network remodeling underlying persistent hyperexcitability, has hindered progress in identifying common mechanistic pathways. ^70–72^

An important limitation of previous studies with respect to standardization relate to recording and analysis methods and dosing variability, using a range of toxin concentrations spanning two orders of magnitude ^10,12,13^. For instance, in a rat model of acquired epilepsy, injections of 15 ng of TeNT in visual cortex caused chronic generalized convulsive seizures^13^. Brener et al injected 0.2 – 3 ng in rat parietal cortex to elicit permanent neuronal synchronization^32^. In mice it was shown that 0.1-0.2 ng TeNT injected in layer 5 of primary visual cortex causes transient reduction in synaptobrevin-2 expression in both inhibitory and excitatory neuronal subtypes and sustained electrographic discharge activity over 60 days post injection ^12^. However, neural activity in that study was assessed primarily with LFP recordings of limited spatial resolution, and a rigorous separation of ictal from interictal activity was not performed. Notably, the reported “ictal” discharges lacked overt motor or behavioral correlates and therefore did not correspond to generalized tonic–clonic seizures; moreover, the electrographic phenotype of these events was not characterized in detail. These methodological differences have complicated efforts to extract common mechanistic principles of epileptogenesis.

Here, we introduce a multimodal, chronic in vivo framework that combines EEG, widefield and two-photon calcium imaging, and awake single-cell patch-clamp recordings to study the emergence of focal epileptiform activity following a single intracortical TeNT injection. Our approach is the first of its kind and allowed us to bridge spatial and temporal scales—from membrane potential dynamics to population-wide activity—revealing the gradual development of *optically defined cortical microseizures* that evolve over weeks and are largely invisible to conventional surface EEG recordings. Local hypersynchrony has been described in rodents exposed to convulsant agents such as pilocarpine, kainic acid, and 4-aminopyridine^36,52–55^, and intracranial recordings in patients with epilepsy have revealed isolated cortical microseizures with spatiotemporally restricted footprints^56,57^ comparable to the microseizure events we report here. While these studies have been instrumental in defining key features of hypersynchrony across systems, they have not delineated the progressive course of focal epileptogenesis at the neural-ensemble level using a variety of complementary, chronic in vivo recordings modalities – a gap addressed by the present study. Our EEG recordings revealed a delayed emergence of spike-and-wave discharges (SWDs^41^ ; fig. 2A) and isolated interictal spikes (fig. 2A/B), beginning approximately three weeks after TeNT injection. On the face of it, our event counts differ from the only previously published work studying focal TeNT in mouse visual cortex, which reported “ictal” epileptiform events at up to 80 events/hour (compared to our ∼3-10 spikes and SWD’s per hour), 10-45 days after TeNT injection^12^. This difference may be explained by three factors: 1) they used more invasive LFP electrodes, whereas we used surface electrodes, 2) the events they classified as seizures did not exhibit “a clear motor/behavioral correlate,” and 3) their definition of epileptiform events did not correspond to our more restrictive categorization of spike-wave discharges and interictal spikes (however we did see frequent uncategorized events without a clear frequency- or rhythmicity related signature).

Interictal spikes and SWDs did not occur in sufficient numbers across all time periods to quantify neuronal engagement with these EEG patterns. However, for the uncategorized EEG events, individual neurons could be classified^38^ as transiently activated or suppressed during these events, but their overall engagement did not increase systematically over time (Suppl. Fig. 2D). These findings are consistent with prior observations that many EEG-defined epileptiform patterns reflect large-scale synchronization without proportional recruitment of local neuronal populations^73,74^.

In contrast, chronic calcium imaging uncovered the gradual appearance of high-intensity, spatially confined microseizure events, whose spatial extent closely matched the original TeNT injection zone and whose neuronal recruitment was strikingly high: 49-83% of neurons in L2/3 versus 32-52% in L4 were activated, while none were suppressed (see Fig. 4). Thus, during peak optically recorded microseizure epochs, microseizures engaged the majority of neurons within the affected field of view in both layers 2/3 and 4, with little evidence of concurrent suppression. Interestingly, these events were essentially undetectable by EEG electrodes placed 2-4 mm millimeters away from the TeNT focus (Suppl. Fig. 2A-C), underscoring the spatially restricted nature of the underlying pathology and highlighting a fundamental limitation of macroscale electrophysiological monitoring for detecting focal epileptogenesis. Even relatively large (i.e. ∼1mm radius, Fig. 3) microseizures are apparently lost to surface electrodes 2-4 mm away, highlighting that optical methods will be necessary to dissect in detail the circuit mechanisms of focal epileptogenesis.

The temporal evolution of microseizures further supports their relevance as a core pathological feature. We followed the evolution of microseizure events using chronic two-photon imaging, and found that they emerged within the first three weeks after TeNT injection (Fig. 4), peaked in numbers and duration around day 30 after TeNT injection, and then gradually declined over subsequent weeks, in parallel with—but not identical to—the time course of EEG abnormalities. We observed microseizures both in layers 2/3 and 4. Layer-specific analyses revealed subtle differences: microseizures appeared slightly earlier in layer 4, whereas the fraction of recruited neurons continued to increase in layer 2/3 but fell in layer 4 during the peak period, suggesting lamina-specific compensatory plasticity mechanisms^75,76^. These dynamics are consistent with prior reports of layer-dependent circuit resilience and adaptation.^77–79^ To obtain a more fine-grained picture of the temporal sequence of neuronal recruitment into microseizure events, we employed in vivo single-cell patch clamp recordings. These provided additional mechanistic insight, revealing that most neurons recruited into microseizure events undergo large, sustained membrane depolarizations (fig 5B), reminiscent of previously described paroxysmal depolarization shifts^80,81^. Interestingly, the cellular calcium fluorescence signal reflected both AP production and AP-free depolarization. It reached the threshold set for robust microseizure detection a full ∼1 sec after neuropil calcium and subthreshold membrane voltage rose significantly (Fig. 5C-H), indicating that microseizures are preceded by a build-up of synaptic and subthreshold network activity. The delay between neuropil activation and robust somatic calcium signals highlights the importance of subthreshold processes in seizure initiation and underscores the limitations imposed by the temporal kinetics of calcium indicators. Future voltage imaging approaches will be critical for resolving these dynamics with higher fidelity, and will allow us to quantify both excitatory and inhibitory synaptic input, and how this might change dynamically in the lead-up to individual microseizures.

Beyond discrete epileptiform events, we observed substantial reorganization of interictal network activity. We serially imaged layer 2/3 and layer 4 fields of view (FOVs) and analyzed interictal activity leading up to microseizure event emergence and reaching up to and beyond D45 for most recordings (Suppl. Table 1 lists animals and recorded time points.)

Immediately following TeNT injection, overall activity levels, calcium amplitude and synchrony transiently increased, more prominently in layer 2/3 than layer 4 (suppl. Fig 7), consistent with the known early effects of TeNT on presynaptic inhibition^8,10,25,63^. Since TeNT primarily targets GABAergic synapses initially^25,63^, and cortico-cortical inhibitory synaptic density is higher in layer 2/3 than layer 4^82^, this likely explains why L2/3 exhibits more prominent network disruption. Over the ensuing 3 weeks, however, interictal network activity was further disrupted, especially in layer 4 but also in layer 2/3, as microseizures started to emerge. Specifically, in L4, overall mean neuronal dΔF/F activity was reduced significantly, with decreased event rate and AUC (a combined measure of integrated firing rate and burst duration), as well as reduced synchrony (Fig. 6). In layer 2/3, only amplitude and AUC were reduced, while event rate was elevated, together mirroring the immediate effect of less intense inhibition relative to layer 4. Functional connectivity measures were disrupted in both layers, most prominently manifesting by a decrease in the number of clusters of coactive neurons and reduced cluster overlap, indicating a breakdown of normal modular organization with a reduced ability of neurons to form and participate in multiple functional modules. These changes coincided temporally with the emergence of microseizures, suggesting that epileptogenesis is accompanied not by global hyperactivity, but by a reduction of ongoing activity and a destabilization of interictal network structure that permits intermittent runaway synchronization. These observations align with prior work reporting delayed TeNT interference of both GABAergic and glutamatergic synaptic transmission^83,84^: healthy cortical circuits operate within a constrained dynamic range, avoiding both excessive quiescence and uncontrolled excitation, whereas, according to our data, TeNT appears to progressively expand this range locally: interictal activity becomes suppressed and decorrelated, while the network remains vulnerable to abrupt transitions into highly synchronized microseizure states. This duality reconciles the apparent paradox of reduced baseline firing coexisting with episodic hypersynchrony.

Microseizures subsided slowly over the next 3.5 weeks (D21-45), and this was accompanied by a normalization of neuronal activity patterns in layer 2/3 (Suppl. Fig 6A), whereas most neuronal activity metrics, except clustering parameters, remained suppressed in layer 4. Interestingly, across D1-20 and D21-45 time bins, neuropil activity was suppressed in layer 2/3, but not in layer 4 (Suppl. Fig 6B). These findings indicate that TeNT alters aggregate synaptic activity—captured by neuropil signals, which reflect summed dendritic and axonal inputs^85,86^—via mechanisms that differ from its effects on cellular spiking and lamina-specific network patterns (Fig. 6). This synaptic disturbance appears to initiate a slow, self-sustaining circuit reorganization^10,87^ that ultimately gives rise to spontaneous epochs of runaway hyper-synchrony. Future studies should leverage higher-temporal-resolution approaches, including in vivo–optimized voltage reporters, to resolve pre-microseizure network dynamics and connectivity changes on the timescale of individual action potentials, thereby defining how normally constrained interictal population activity transitions into epileptiform events.

Interneuron-specific analyses further illuminate the underlying circuit imbalance. Neither PV^+^ nor SST^+^ interneuron number decreased over time (fig. 7A-C), consistent with prior reports suggesting that TeNT injections do not cause anatomical injury to the tissue. However, PV^+^ interneurons exhibited disproportionately low activity relative to surrounding pyramidal neurons (Fig. 7D-E). PV^+^ interneurons exert strong, fast perisomatic inhibition and have been shown in other settings to play an important role in containing run-away activity^88^. Their relatively weak activity then is a symptom of E/I imbalance in favor of excitability in the local cortical microcircuit. PV^+^ cells also play a role in maintaining precise spike timing, particularly during low-drive states^89^. Suppression of PV^+^ interneuron activity can therefore impair synchronization of precise spike times across multiple neurons when population firing rates are low^89^ – which is typically the case when the animal is in a state of quiet wakefulness with no incoming sensory stimulus, like in our experimental setup - and is thus consistent with our observation of reduced network synchronization in layers 2/3 and 4 interictally during the time period of increasing microseizure emergence (day1-20, figure 6). In contrast, SST^+^ interneurons were found to be relatively hyperactive. SST^+^ cells exert strong inhibitory effects on pyramidal neuron dendrites^65^, shaping how they integrate inputs and undergo plasticity and driving feedback inhibition to the local population^89^, but their ability to directly suppress pyramidal neuron spiking is typically weaker than that of PV^+^ interneurons^66^. While SST^+^ cells primarily target dendrites and modulate input integration rather than directly suppress spiking, their increased activity may reflect a local compensatory attempt to restore excitatory–inhibitory balance. This may succeed in compensating most of the time, yet fail to do so during microseizure events. In the single-cell patch-clamp data we observed an apparent dip in the overall activity of pyramidal neurons immediately prior to the microseizure events (figure 5E; this was not visible in calcium imaging –Suppl. Fig. 8--, likely due to the slow kinetics of the calcium reporter), suggesting a transient epoch of increased inhibition occurs immediately preceding the onset of microseizures. Notably, this occurs despite the relative suppression of PV^+^ cell activity we observed, and may be mediated by increased inhibition SST^+^ interneurons exert on pyramidal cells. However, it is well known that SST^+^ interneurons directly inhibit PV^+^ cells^67^, raising the possibility that, when SST^+^ interneurons become hyperactive, they may secondarily, e.g. after a certain delay, directly and strongly suppress PV^+^ interneurons, eliminating the residual PV^+^ control of pyramidal cells and thereby facilitating the transition into microseizures. Future imaging studies using reporters with greater sensitivity and better temporal resolution to simultaneously image multiple interneuron subpopulations, should be able to resolve the interactions leading to microseizure emergence.

In summary, by combining chronic optical imaging with electrophysiological recordings across scales, we characterized the minimally invasive TeNT mouse model of focal epileptogenesis, revealing how lamina-specific circuit reorganization gives rise to spatially restricted microseizures undetectable by conventional surface EEG. Our findings emphasize that focal epilepsy can emerge from progressive interictal circuit destabilization, rather than sustained hyperactivity, and point to differential roles of PV^+^ and SST^+^ interneurons in shaping this trajectory. As next-generation reporters for voltage, neurotransmitters, and neuromodulators mature, this framework will enable deeper dissection of the mechanisms by which local excitatory–inhibitory balance breaks down, how microseizures emerge and propagate, and how focal pathology may generalize through thalamocortical networks.

## Supplementary material

**Supplementary figure 1 (supplement to figure 2):**
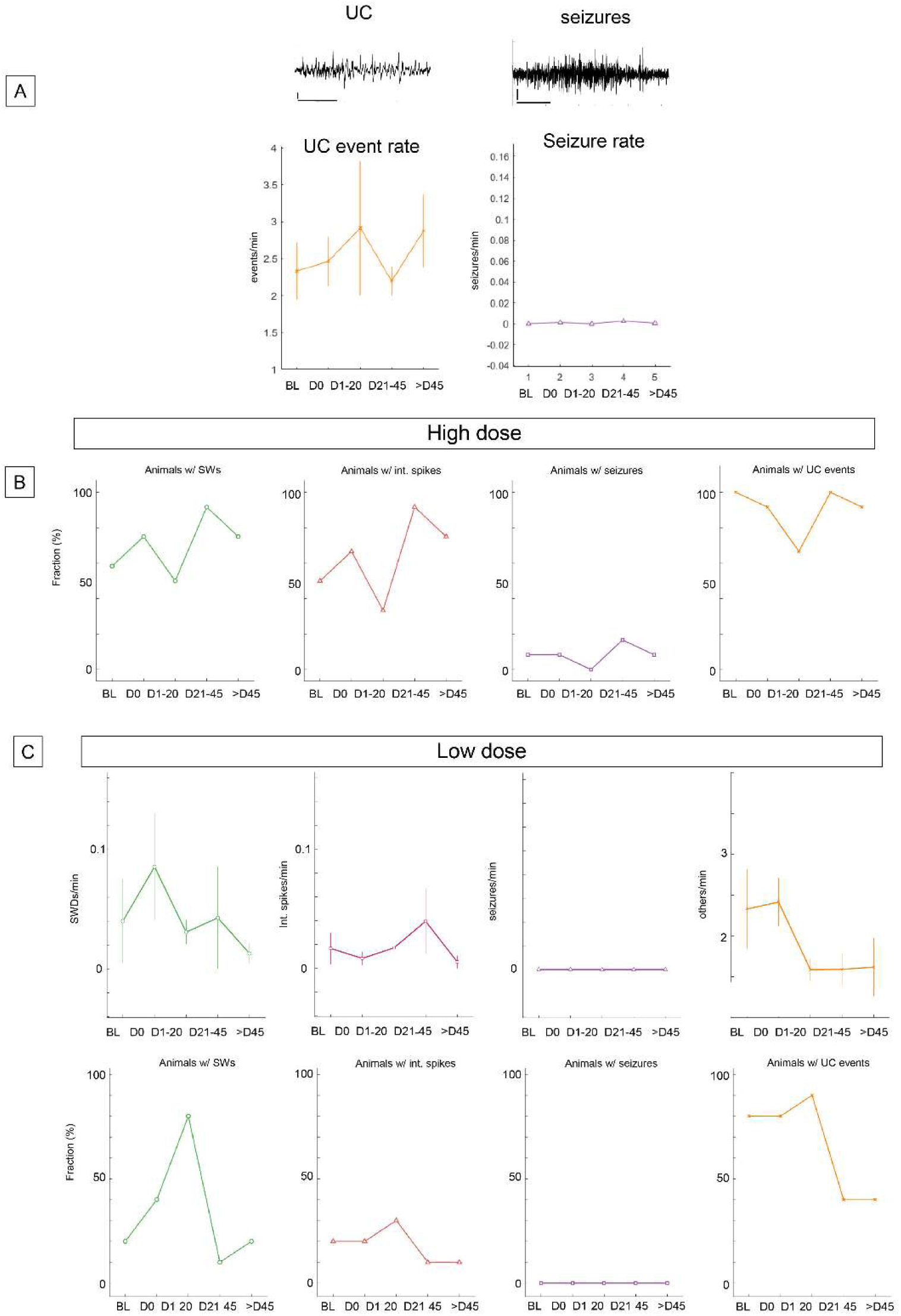
additional EEG analysis quantification **A:** Top: Examples of uncategorized (UC) EEG event, and a generalized electrographic seizure, bottom: quantification of the rates of these types of events measured across the time bins BL (baseline), D0 (immediately after TeNT injection, D1-20, D21-45, and >D45. Uncategorized events were not significantly more frequent after TeNT application, indicating that these events may also occur naturally, and are therefore not a relevant marker of TeNT-induced network remodeling. Seizures were so rare that we could not analyze their rates and characteristics further in a meaningful way. **B:** Fraction of animals that after TeNT injection developed (from left to right) uncategorized events, interictal spikes, seizures, and SWDs. **C:** Quantification of the rate of the 4 different EEG event categories (top) and the fraction of animals that developed them, upon low-dose TeNT application.

**Supplementary figure 2, supplement to figs 3,4:**
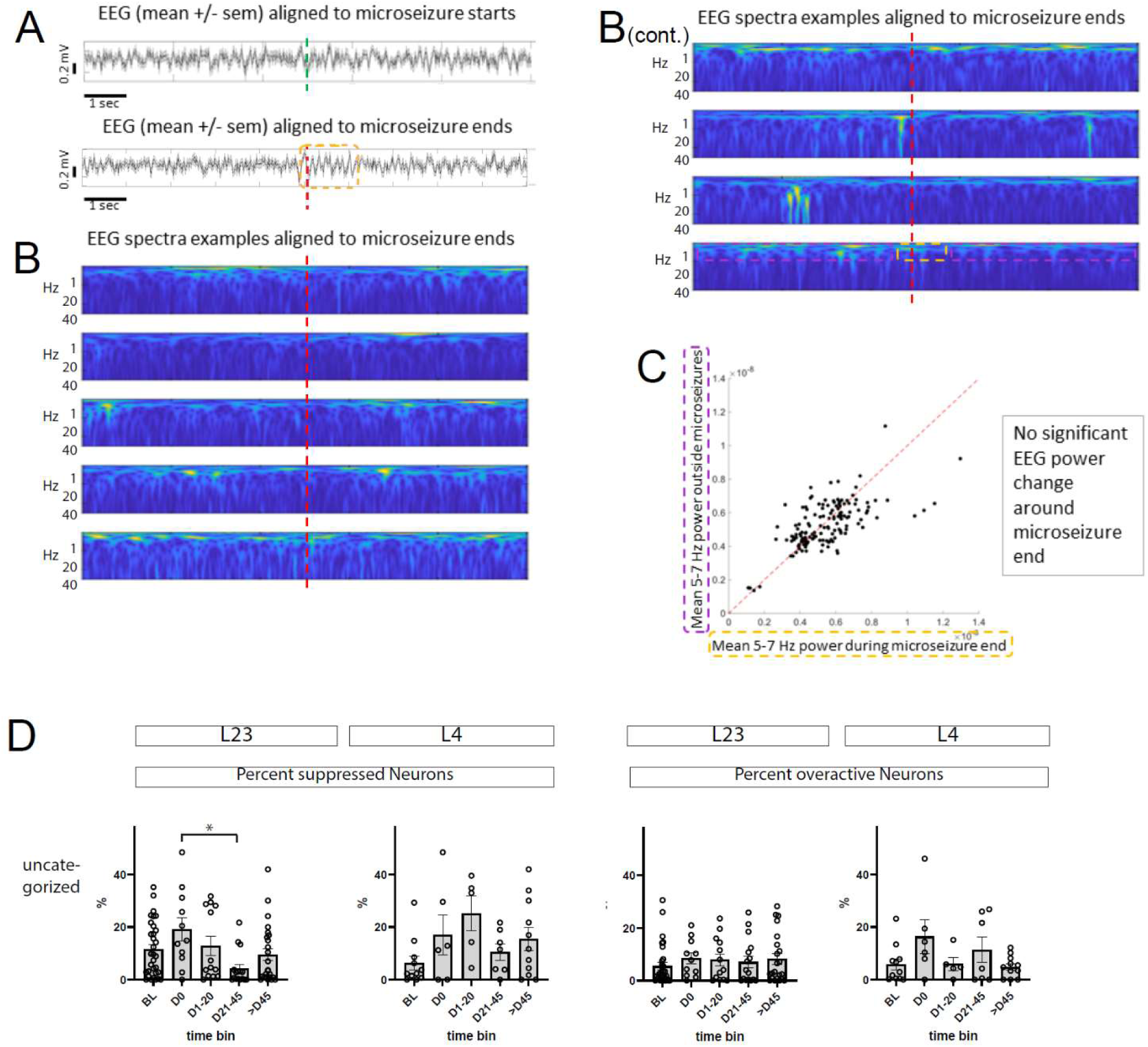
Reverse correlation of EEG traces to calcium microseizure start/end times shows no significant correlation; Neuronal responses during abnormal EEG discharges were not significantly modulated as a function of time after TeNT injection: **A:** Reverse correlation of EEG traces to calcium signal derived microseizure start and end times, showing no significant correlations. A: EEG (mean±s.e.m., shaded grey error bars), aligned to microseizure start time (green dashed vertical bar, top) and end time (red dashed vertical bar, bottom). While there was no appreciable correlation between EEG and microseizure start time points, visual inspection revealed some low-frequency activity around end points, which is why we analyzed those time points further below. **B:** examples of EEG spectra (1-40 Hz) from randomly selected microseizures, aligned to end time points. **C:** scatter plot of mean EEG signal (5-7 Hz band pass filtered) around microseizure end (x-axis) vs. interictal time periods (y-axis), showing no significant increase or decrease in low-frequency EEG power around microseizure end time points. **D:** Neuronal activity during EEG events: In a portion of imaged neurons, cellular calcium activity was elevated during these events, in others suppressed. This analysis was consistently possible only for UC events (SWD and interictal spike events too rare and/or short in most recordings for ranksum comparison of dΔF/F concatenated traces).

**Supplementary figure 3, supplement to figure 4:**
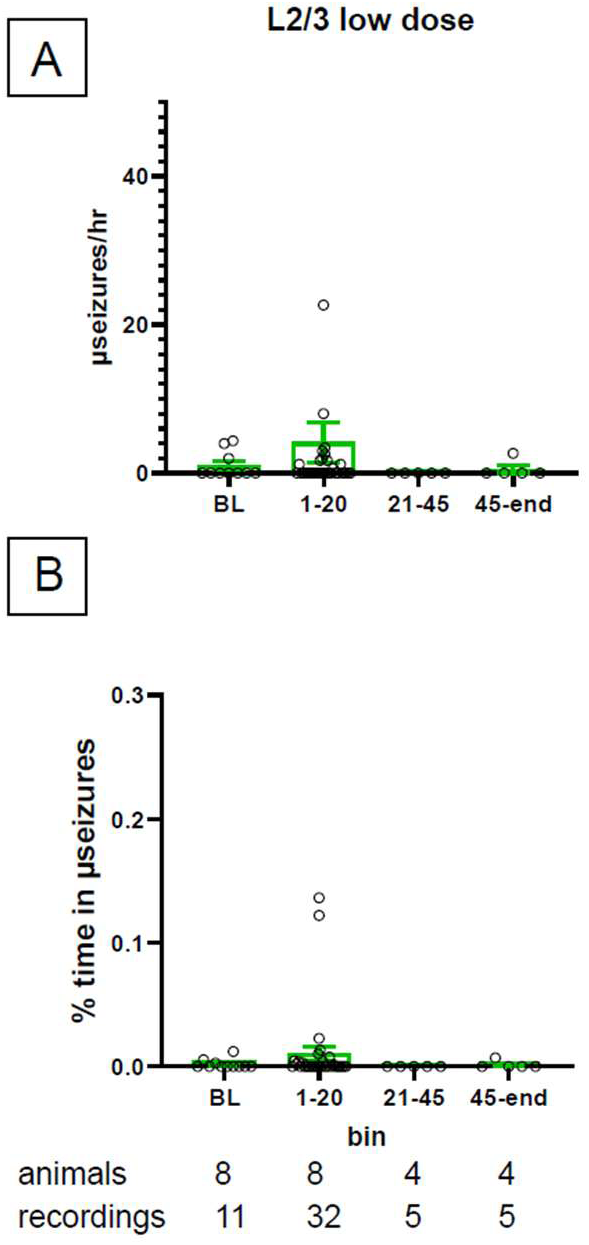
low-dose TeNT effects. **A:** quantification of microseizure rates recorded before and after low-dose TeNT injection, using the same time bins as in figure 4. These recordings were only made in layer 2/3 and show a small number of microseizures in the early (D1-20) time bin, which in contrast to the high-dose results, do not develop into more frequent events after D21. **B:** similarly to A, the percentage of time in microseizures per recording did not exceed baseline levels in the D21-45 time bin.

**Supplementary figure 4, supplement to figure 4:**
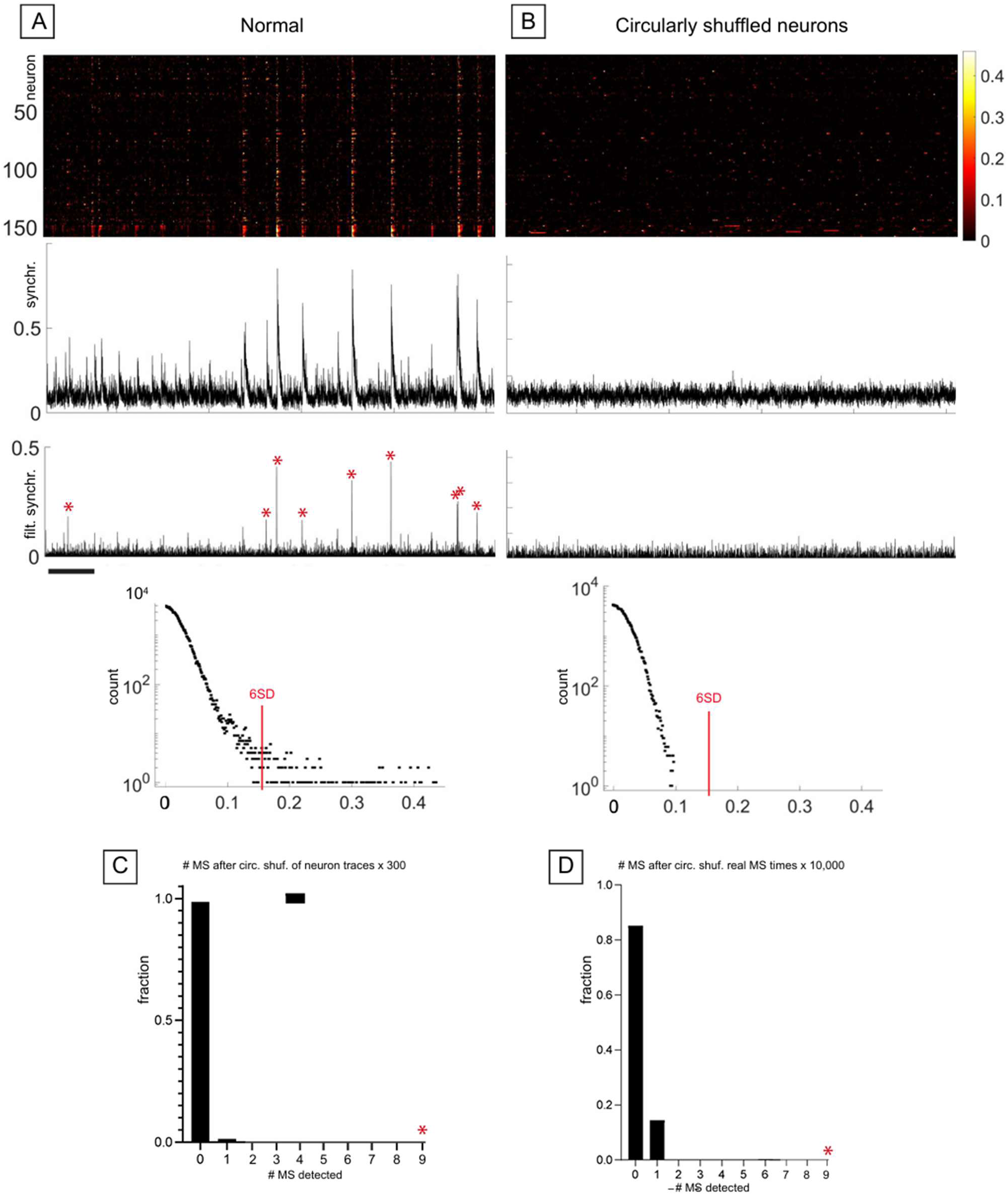
Validation of microseizure detection, example animal with shuffled distributions. **A/B:** ΔF/F calcium raster plot of an example recording with clear microseizures visible (left), and after scrambling neuronal calcium traces via circular shuffling by random offsets. Below is the synchrony (i.e. number of coactive ROIs across time) for the normal and scrambled versions, as well as a bandpass-filtered version of the synchrony which is thresholded at 6SD to identify microseizures (red stars). Next (below) we show semi-logarithmic histograms of the raw (unfiltered) synchrony and the 6SD cutoff (as applied to the filtered synchrony), showing plenty of candidate microseizures for the normal version (many of which get removed after filtering), and no candidates for the scrambled version. **C:** number of microseizures detected after repeating the circular shuffling of neuron traces 300 times, showing that in most (>98%) of repetitions, zero microseizures were detected. **D:** Using an alternative scrambling method by moving the detected microseizure times by a random amount of time and computing the number of overlapping scrambled microseizures with the actual ones, we find that the true number of microseizures was impossible to reach for the scrambled versions after 10,000 iterations, further confirming the validity of our microseizure detection approach.

**Supplementary figure 5:**
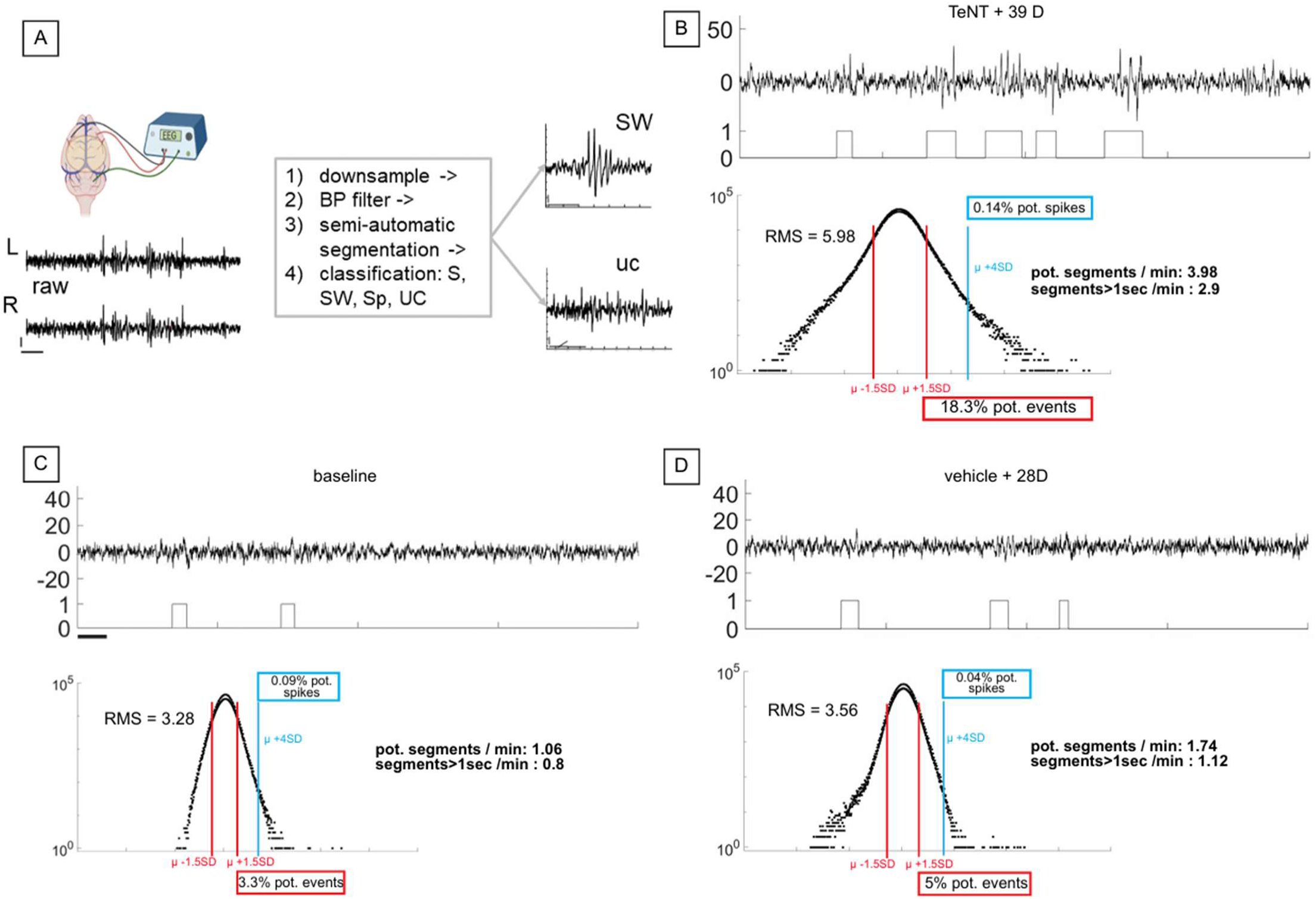
example of EEG event identification and noise distribution, example animal with TeNT, vehicle, and control activity. **A)** 2-channel EEG (ref over cerebellum) was recorded at 10kHz, and subsequently re-referenced, downsampled to 100Hz, bandpass-filtered, before semi-automatically finding segments of interest and classifying them manually (example of a spike-wave discharge (SWd) and an uncategorized high-amplitude event shown here, horizontal bars = 1 sec, vertical bars = 0.1 mV). **B)**Top: Band-pass filtered EEG example (20 sec) and the result of the thresholding algorithm providing time periods of potential EEG discharge events which were then classified by visual inspection. Bottom: semi-logarithmic histogram showing thresholds for potential abnormal events (red vertical lines) and interictal spikes (blue lines). **C)** Analogous to B, EEG from a baseline recording before vehicle injection yields a significantly lower number of potentially abnormal EEG patterns identified by the thresholding algorithm. **D)** Analogous to C, EEG recorded 28 days after vehicle injection did not develop additional EEG discharge patterns as observed after TeNT injection (A).

**Supplementary figure 6 (supplement to fig 6):**
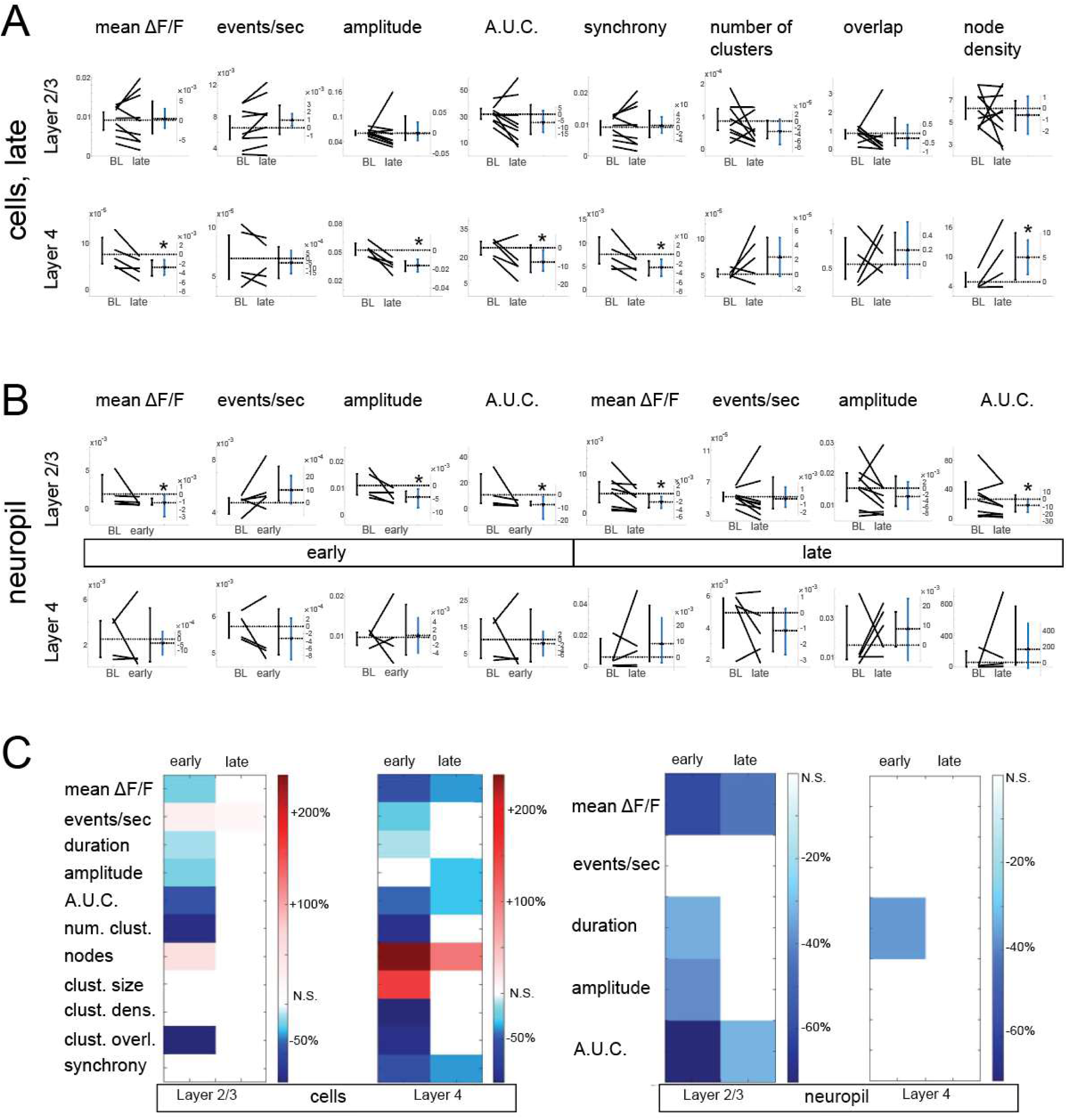
Late calcium activity changes upon TeNT injection, neuropil effects, summary of effects. **A:** Comparison of cellular activity metrics between baseline (BL) and the D21-45 time bin using estimation statistics, similar to figure 6. Paired tests between baseline (BL) and late (D21-45) time bins are shown, as well as the mean and 95% CI (blue vertical bar) of the differences. Significance was established when the 95% CI did not overlap with the zero difference (horizontal black dashed line corresponding to the mean of the BL distribution, intersecting the zero on the secondary (delta) y-axis; the other horizontal dashed line corresponds to the mean of the other (late/early) distribution) **B:** comparison of neuropil activity metrics between baseline and D1-20 data (left half) and baseline vs. D21-45 (right half). Estimation statistics were used, analogous to A). Clustering-based metrics were not computed as they require cellular signals. **C:** Heatmaps summarizing activity metric comparisons across layers, time bins, and cells/neuropil, color coded as percent change versus baseline.

**Supplementary figure 7 (supplement to fig 6):**
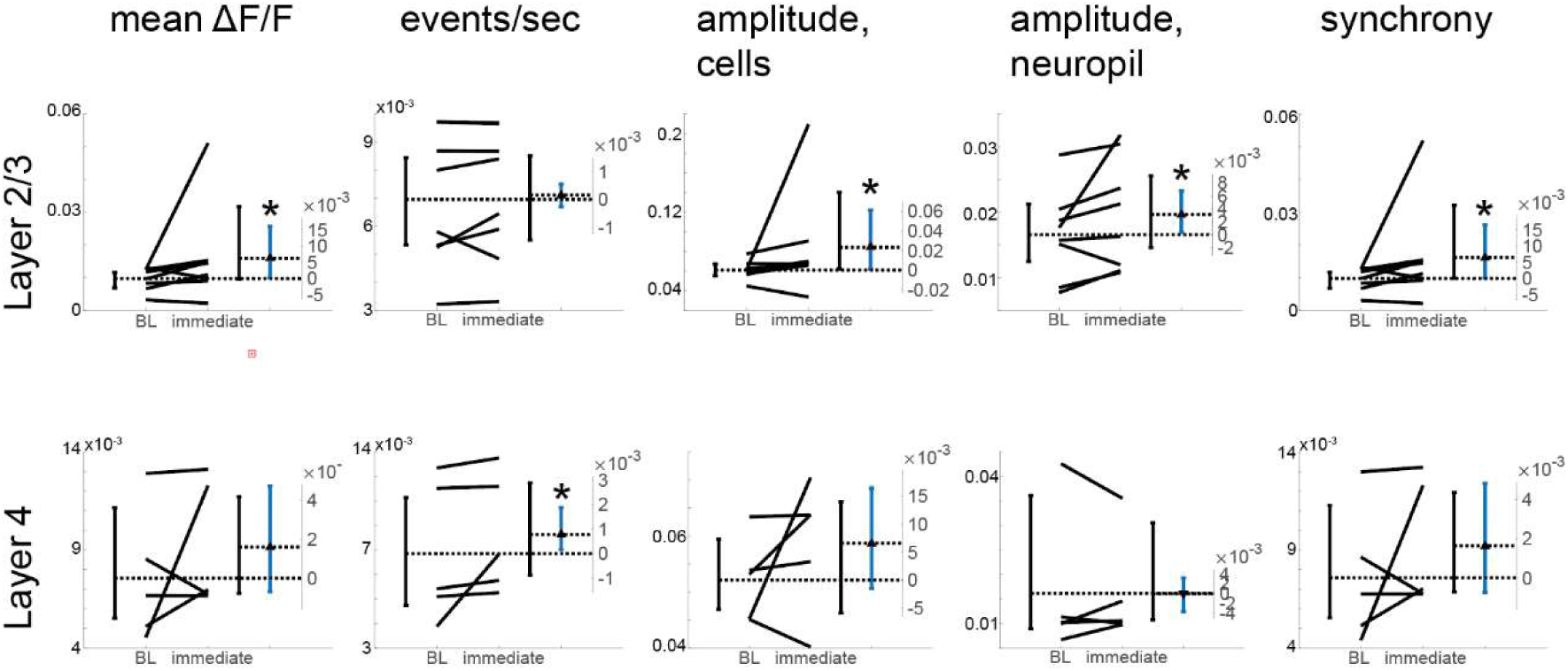
Immediate cellular calcium activity changes upon TeNT injection. Immediate effects of TeNT injection, comparing baseline activity recorded immediately prior to injection with activity recorded within 2 hours of injection. Within this time period, most activity metrics in layer 2/3 as well as event rate in layer 4, were elevated. Estimation statistics were applied analogously to suppl. fig. 6.

**Supplementary figure 8 (supplement to figure 6):**
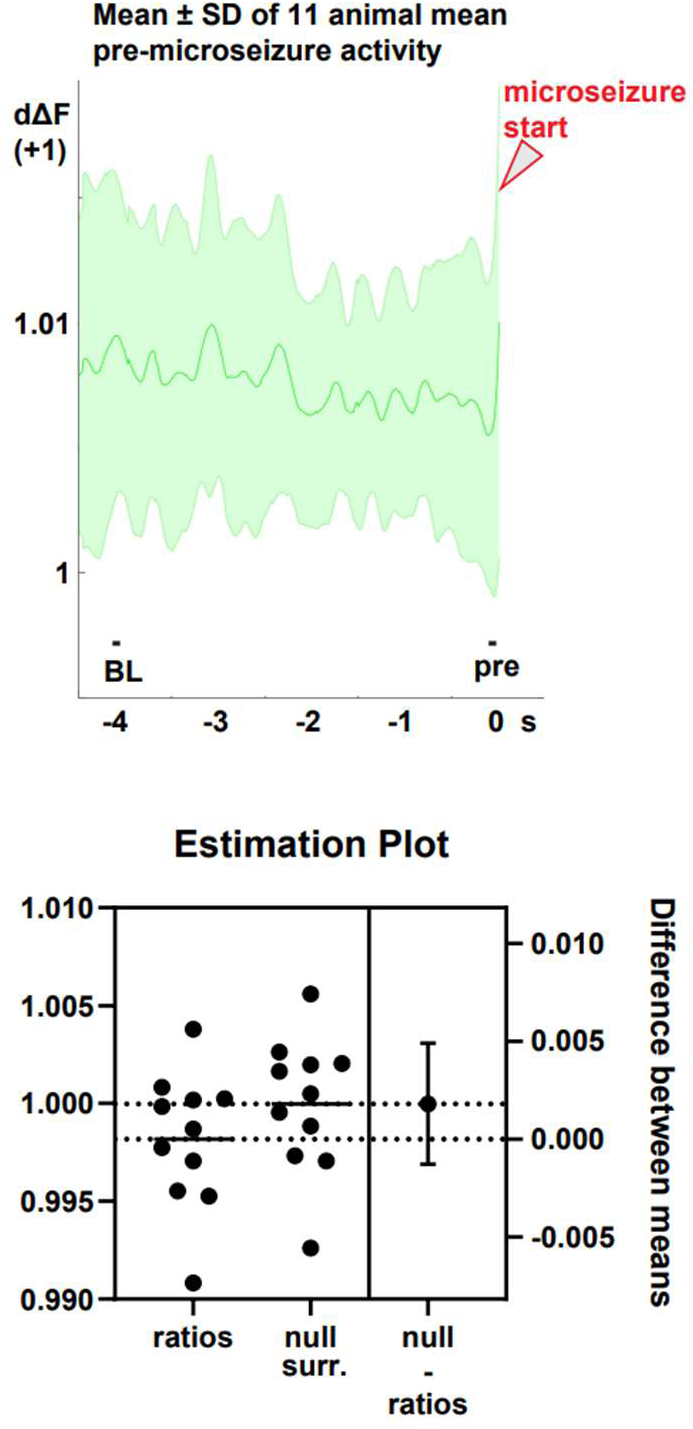
**Top**: Deconvolved DF/F activity (4.5 seconds preceding microseizure onset times) was extracted from 11 animals (L2/3 recordings only). The traces from each animal were averaged, and here we plot the mean +/- SD of those averages. **Bottom**: for each animal we calculated the mean pre-microseizure activity level (50ms immediately prior to microseizure onset), and the mean baseline (BL) value for comparison. For each animal we calculated the ratio of pre/BL values. The mean ratio was below 1 (exact value = 0.9982). To test for significance, we shifted the distribution of measured ratio up to match a mean of 1 (which would be expected if there were no effect). We compared this “null surrogate” distribution with the actual ratios and found no significance (p = 0.24, unpaired t-test).

**Supplementary table 1:**
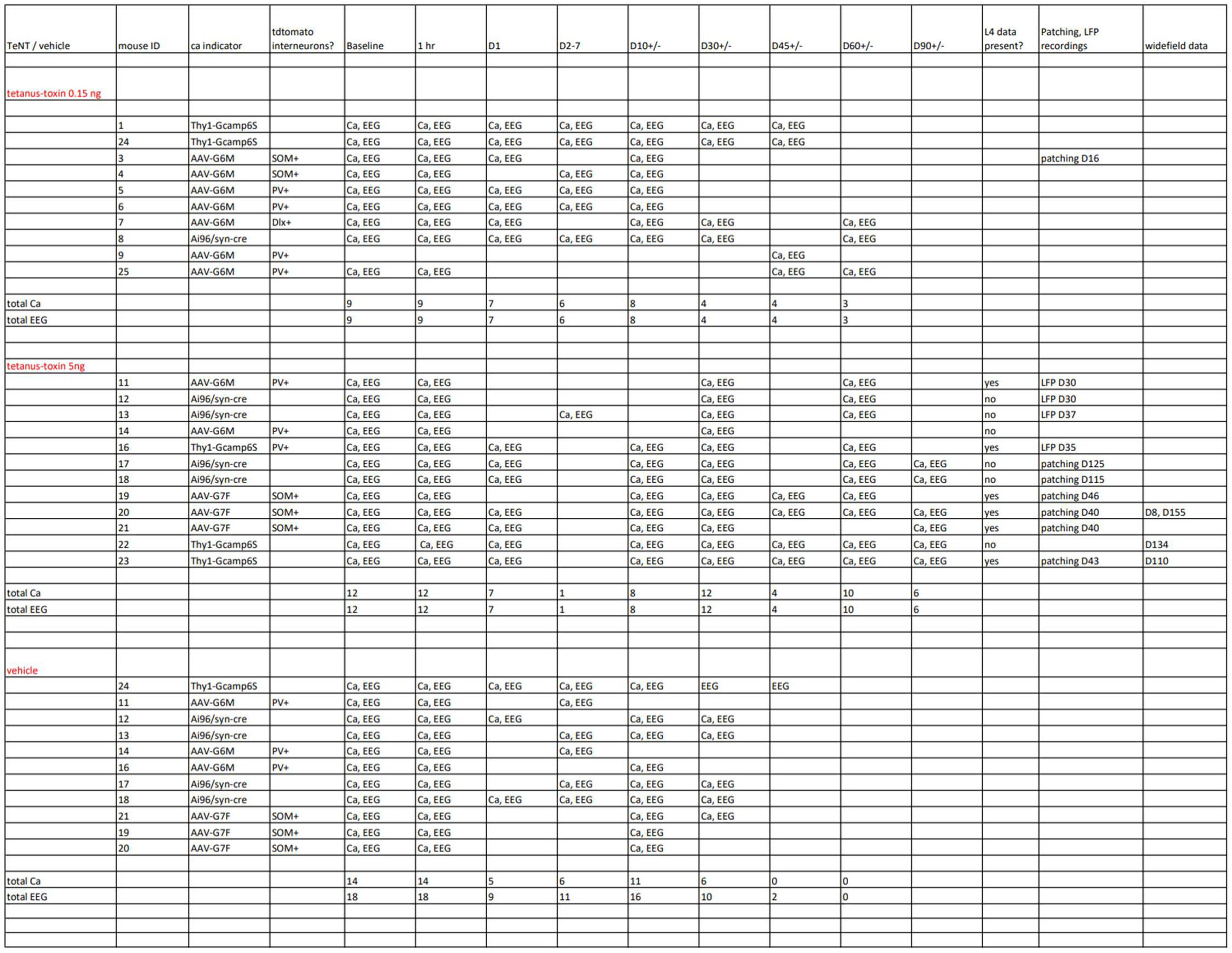
all animals used for this study with data types and time points recorded.

